# cOmicsArt – a customizable Omics Analysis and reporting tool

**DOI:** 10.1101/2024.10.30.621025

**Authors:** Lea Seep, Paul Jonas Jost, Clivia Lisowski, Hao Huang, Stephan Grein, Hildigunnur Hermannsdottir, Katharina Kuellmer, Tobias Fromme, Martin Klingenspor, Elvira Mass, Christian Kurts, Jan Hasenauer

**Affiliations:** Computational Biology, Life & Medical Sciences (LIMES) Institute, University of Bonn, 53115 Bonn, Germany; Institute of Molecular Medicine and Experimental Immunology (IMMEI), University Hospital of Bonn, University of Bonn, 53127 Bonn, Germany; Developmental Biology of the Immune System, Life & Medical Sciences (LIMES) Institute, University of Bonn, 53115 Bonn, Germany; Chair of Molecular Nutritional Medicine, TUM School of Life Sciences, Technical University of Munich, Freising, Germany; EKFZ—Else Kröner-Fresenius Center for Nutritional Medicine, Technical University of Munich, 85354 Freising, Germany; ZIEL Institute for Food & Health, Technical University of Munich, 85354 Freising, Germany; Helmholtz Center Munich, German Research Center for Environmental Health, Computational Health Center, 85764 Neuherberg, Germany

## Abstract

**Motivation:** The availability of bulk-omic data is steadily increasing, necessitating collaborative efforts between experimental and computational researchers. While software tools with graphical user interfaces (GUIs) enable rapid and interactive data assessment, they are limited to pre-implemented methods, often requiring transitions to custom code for further adjustments. However, most available tools lack GUI-independent reproducibility such as direct integration with R, resulting in very limited support for transition.

**Results:** We introduce the customizable Omics Analysis and reporting tool – cOmicsArt. cOmicsArt aims to enhance collaboration through integration of GUI-based analysis with R. The GUI allows researchers to perform user-friendly exploratory and statistical analyses with interactive visualizations and automatic documentation. Downloadable R scripts and results ensure reproducibility and seamless integration with R, supporting both novice and experienced programmers by enabling easy customizations and serving as a foundation for more advanced analyses. This versatility also allows for usage in educational settings guiding students from GUI-based analysis to R Code.

**Availability:** cOmicsArt is freely available at https://shiny.iaas.uni-bonn.de/cOmicsArt/

User documentation is available at https://icb-dcm.github.io/cOmicsArt/

Source code is available on GitHub https://github.com/ICB-DCM/cOmicsArt

A docker image can be retrieved from https://hub.docker.com/r/pauljonasjost/comicsart/tags

A snapshot upon publication can be found on Zenodo: https://zenodo.org/records/13740904

A screen recording of cOmicsArt is available at: https://www.youtube.com/watch?v=pTGjtIYQOakp

**Supplementary information:** Supplementary data are available online.

## 1 Introduction

The generation of vast amounts of bulk-omics data has become a common practice. The data analysis often greatly benefits from collaborative efforts between experimental (data generator) and computational (data analyst) researchers as complementary expertise is needed in bioinformatics practices (Attwood *et al*. 2019). Particularly during the initial stages of analysis, broad explorations, comparison to domain knowledge, and statistical analysis are crucial. The early phase of the analysis allows for data-driven successive refinements (Cathy O’Neil and Rachel Schutt 2013), where tabular results and visualizations serve as common ground for the collaborators exchanging ideas, hypotheses, and knowledge, ultimately guiding the further, more tailored analysis.

Many software tools with graphical user interfaces (GUIs) have been developed to support bioinformatics practices, offering a range of analysis levels from basic to advanced, for both, specific datasets and general use cases. GUIs are particularly valuable in bioinformatics as they make data analysis more accessible to users with varying levels of expertise, and enable rapid, often interactive, assessments of data and results. There are also big platforms, such as Galaxy (Abueg *et al*. 2024) and Partek (Partek 2024), which integrate a wide array of analytical tools without the need for programming expertise (for a systematic assessment of available tools see Supplementary Note A–B).

A key challenge in omics data analysis is bridging the gap between GUI-based and custom analysis. GUIs are limited to pre-implemented analyses, often necessitating a shift to custom code for out-of-scope tasks. Many GUIs lack easy reproducibility outside their interface, forcing users to manually redo analyses to extend initial work. Yet, relying solely on custom code sacrifices the advantages GUIs provide, particularly for non-programmers.

Herein, we present a customizable Omics Analysis and reporting tool – cOmicsArt – that combines the advantages of GUIs for the explorative phase with interface-independent reproducibility, enabling swift transition to custom(ized) analysis. cOmicsArt allows users to easily perform tasks such as cluster-, correlation-, principal component-, set-, enrichment- and statistical analysis associated with bulk-omics data and interpret results interactively with inbuilt visualizations (Fig. 1A). With auto-generated reports, R scripts, and convenient data objects, cOmicsArt provides both human- and machine-readable output for all analyses. The cOmicsArt-generated R code that reproduces all GUI-based analyses enables a seamless transition to custom(ized) analyses (Fig. 1B).

**Fig. 1.**
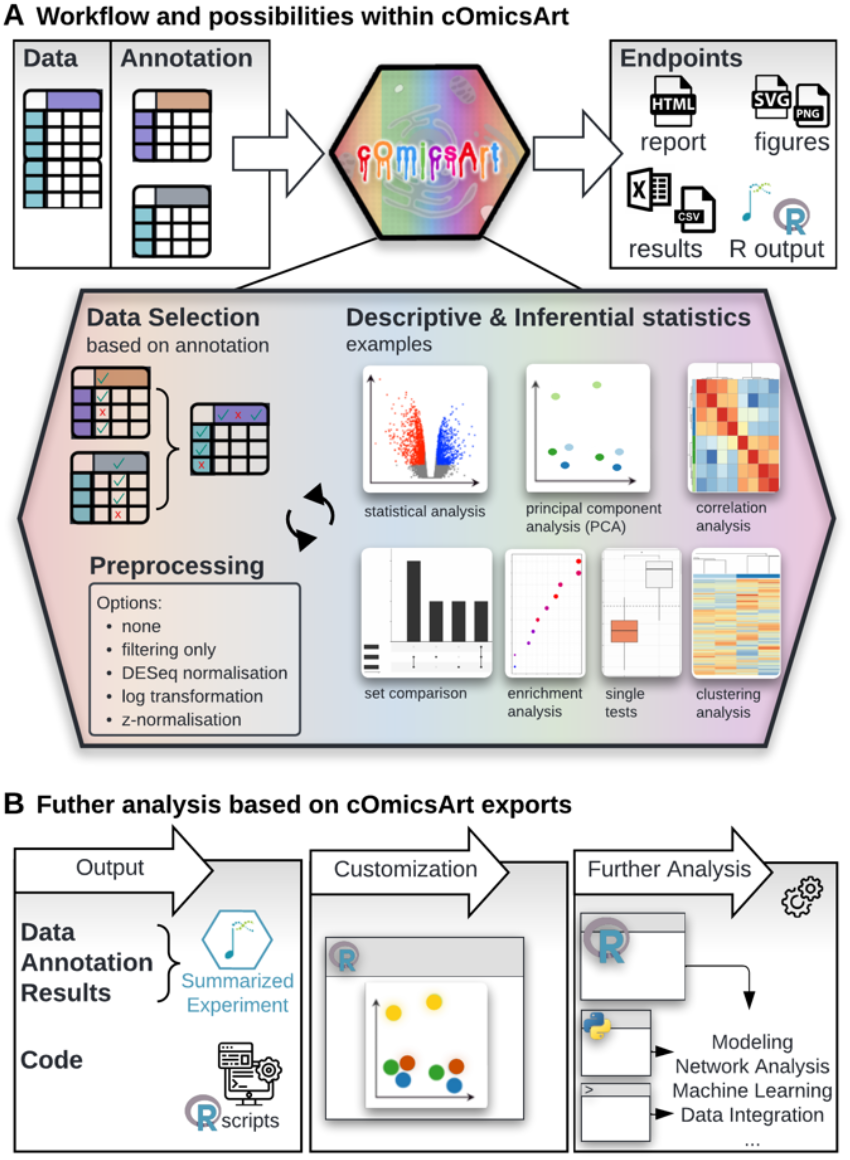
Workflow within cOmicsArt and beyond. **A** cOmicsArt expects the data matrix with samples as columns and entities, e.g., genes or lipids, as rows. Two additional required matrices hold annotation information on the samples and entities respectively. Users can first select data based on provided annotation, and choose from pre-processing options. For the explorative analysis, cOmicsArt offers a variety of modules, such as sample correlation, PCA, statistical testing, enrichment analysis, and different visualization routines. Results can be exported as visualizations and tables. The entire workflow is documented automatically and provided via an HTML report. **B** An additional export option provides the input data accompanied by produced results and user-chosen parameters as a data object as well as the code base to fully reproduce the results and visualizations thereof within R. This output allows to apply advanced customization and can serve as a basis for further analysis.

## 2 Features

### 2.1 User interface and analysis features within cOmicsArt

The cOmicsArt user interface is designed for ease of use, offering quick familiarization and a low entry barrier through a consistent design and progressive expansion of analysis options. The interface includes a data selection tab, multiple analysis tabs, and links to the human-readable reproducibility report and the complete documentation (Fig S1).

Upon launching cOmicsArt, only the data upload tab is available. This tab expects three dataframes: (i) a data matrix with entities (genes, lipids, metabolites) in rows and samples in columns (Fig. S2A), (ii) a sample annotation matrix with additional features as columns (e.g., health status) (Fig. S2B), and (iii) an entity annotation matrix also with additional features as columns (e.g., common names) (Fig. S2C). Alternative options for upload are the replacement of the sample annotation matrix with a so-called Metadatasheet (Seep *et al*. 2024) or the upload of a compressed data object created by cOmicsArt itself upon request. To address difficulties during data upload, we provide extensive guidelines, and an upload check module highlighting potential issues of the data format.

Once data is successfully uploaded, the user is guided to the pre-processing tab - the first step in the analysis. As all analysis tabs, it consists of a side panel for displaying analysis options and a main panel for presenting results, reporting, and download options. Each side panel has two sections: the upper section includes options that trigger re-computation with a “Do/Get Analysis” button, while the lower section adjusts the result appearance in real-time without re-computation. The main panel allows users to interact with visualizations, such as hovering over data points for more information or zooming into specific regions, enabling efficient retrieval of details.

cOmicsArt offers a wide range of features for explorative and statistical analysis upon extensive data selection as well as pre-processing possibilities and stands out compared to similar tools (Table 1). Data can be selected based on the provided annotation, for both samples and entities. Pre-processing includes several options as well as batch correction.

**Table 1.**
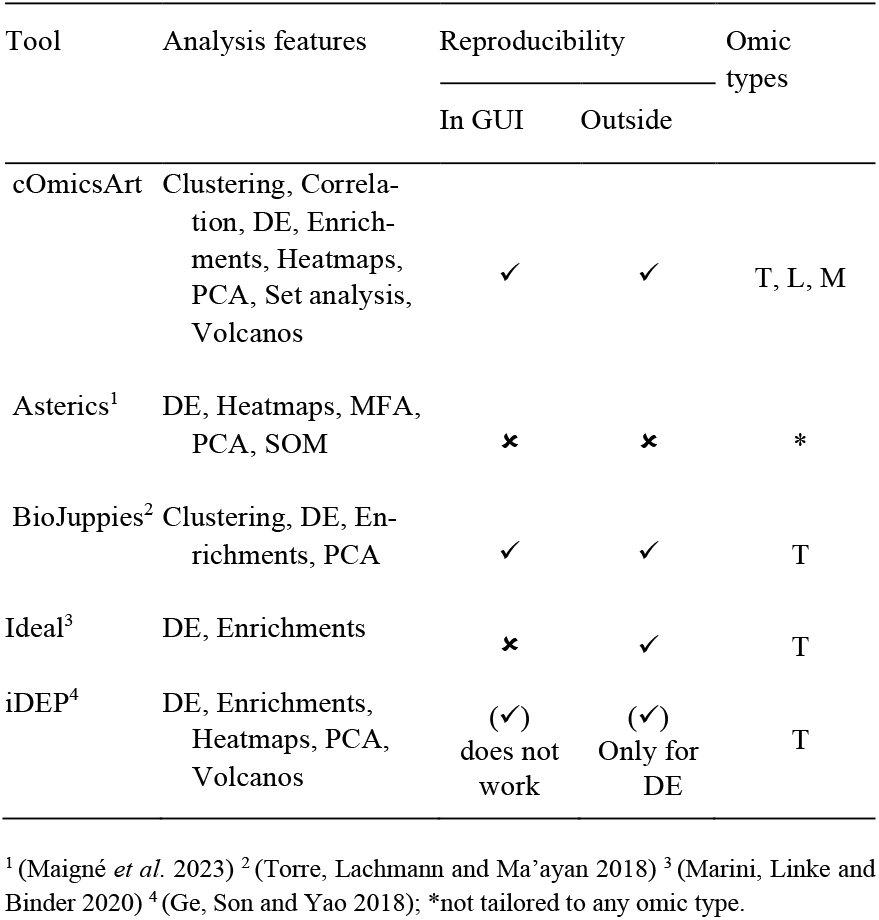
Comparison of a subset of available bulk-omics analysis tools with a graphical user interface. ✓ and ⨯ indicate the absence or presence of respective characteristics. Note, that this is only a subset of identified tools (see Supplementary Table S1 and Supplementary Notes A - B). Abbreviations: DE = differential expression analysis, PCA = principal component analysis, MFA = multi-factor analysis, SOM = self-organizing maps, T = Transcriptomics, L = Lipidomics, M = Metabolomics.

#### cOmicsArt

Explorative analysis in cOmicsArt allows users to assess sample correlations through agglomerative clustering and visualize data in reduced dimensions using principal components. Users can evaluate the loadings of each or across multiple principal components to identify key contributing entities. Further, the pre-processed measurements can be displayed in a heatmap format, which includes clustering of entities and samples. For the heatmap, users can additionally subset entities by choosing options such as displaying the top 20 entities based on the calculated log fold change between two conditions. Statistical analyses can be performed on all possible comparisons within a sample’s annotation, with the ability to conduct multiple tests when comparing more than two groups. The included set analysis allows for the comparison of respective outputs to identify, e.g., shared significant entities. The complete set or derived subsets of interest, if mapped to genes, can be subjected to enrichment analysis. For a systematic assessment of available tools based on additional characteristics, see Supplementary Note A–B.

Users have the option to download all visualizations or results from any analysis tab. Furthermore, one can download the auto-generated HTML report and self-sufficient R scripts (see integration with R). To avoid clutter from iterative refinement of the figures, users can select which visualizations to include in the report and optionally add notes. The report captures all information for reproducibility in a human-readable manner and can display the iterative process of data exploration. Additionally, the report includes several publication blueprints describing the procedure and citing all used packages.

### 2.2 Seamless integration with R

The identified gap between the user interface and further analysis was a key focus in designing cOmicsArt. We ensured seamless integration of analyses done within cOmicsArt with R. For each analysis, users can export a self-sufficient R script, a utility script with custom functions, and all associated data with a single click.

The R script includes all steps, from setting the environment (sourcing utility functions and loading needed R packages), over data selection and pre-processing, to the analysis. The script serves as a framework for customizing visualizations and as a base for further out-of-scope analysis. We explicitly chose not to create a cOmicsArt R package. By providing the option to download self-sufficient R scripts, we allow easy manipulation at top level without options being packed within package functions, aiming to reduce complexity.

The obtained data includes a ‘SummarizedExperiment’-object (Morgan *et al*. 2022) with computed results and a parameter list with all user-specified options. The Summarized Experiment object facilitates integration with many workflows and tools from the Bioconductor universe (Huber *et al*. 2015).

## 3 Use cases of cOmicsArt in research and education

cOmicsArt has been successfully integrated into both research and educational settings, demonstrating its versatility and accessibility. We present two distinct showcases based on two biomedical datasets, followed by an overview of the overall benefits of cOmicsArt as reported for various user groups.

The first showcase includes the web-browser based analysis and visualization of transcriptomics data and obtained results from a study investigating how a high-salt diet impairs immune defense against kidney infections by disrupting neutrophil function (Jobin *et al*. 2020). The analysis covers data retrieval, pre-processing, global pattern exploration, statistical analysis, and documentation, offering step-by-step insights on how to perform each using cOmicsArt (Supplementary Note C, Supplementary E). The second showcase is based on a metabolomics dataset from a study on the effects of maternal obesity on offspring liver health (Mass *et al*. 2023). The showcase presents an integrated analysis workflow beginning with cOmicsArt and leading to subsequent customization of results highlighting both the transition to R and in-depth customization options (Supplement Note D), including also an example code snippet provided by the app (Supplementary F). Both showcases exemplify versatility and can be readily used as starting point for one’s own analysis.

In research, cOmicsArt has been utilized by a diverse group of collaborators, including those with varying levels of programming expertise. Usability testing was iteratively conducted through structured unmoderated observational sessions, where new users were observed while interacting with cOmicsArt, with and without specific tasks, followed by a questionnaire and verbal feedback. Taken observations and feedback refined the development process. For individuals without programming skills, the app offers an intuitive interface that enables ad hoc analysis of their data without the need for coding. The interactive features, comprehensive documentation, and diverse options for result downloads are particularly beneficial in these cases. For those with programming experience, cOmicsArt provides the same ease of use while also offering the advantage of full customization through its integration with R. The used questionnaire is linked within the app to allow ongoing and broader user experience evaluation. cOmicsArt’s value is further amplified in collaborative settings, where new results can be quickly assessed by all parties involved, and scripts from previous analyses can be shared for further exploration.

In educational settings, cOmicsArt was introduced to students new to R programming. Students quickly familiarized themselves with cOmicsArt allowing them to perform analysis despite lacking coding experience. Some encountered the need for more refined analyses, at which point they started working with the generated R code. cOmicsArt thus served as an effective teaching tool, not only simplifying complex biomedical data analysis tasks but also encouraging discussions on the limitations of GUIs and the importance of understanding the R code.

## 4 Implementation

cOmicsArt is a web app – accessible by all common browsers – featuring a modular design and is built within the framework of R ‘shiny’ (Chang *et al*. 2024) (list of all used packages in Supplementary Table S2). Modularization of the backend enables the easy extension of the app to new analyses. The data transfer between the modules is facilitated by two session-specific global data-objects holding (i) the original and processed data as well as analysis-specific results and (ii) the user-specified input parameters (accessible to the user upon R code and data download). Each module’s environment has its namespace allowing for clear separation. Extensive failure checks are implemented with ‘tryCatch’-statements to prevent app crashes and data loss.

Code quality was ensured by leveraging GitHub’s collaborative features, including required code reviews by another person. Implemented tests are executed with GitHub actions. All data and results generated while using the web tool are stored on our servers for the duration of the user’s active session, after which it will be deleted. User can install and run cOmicsArt locally, either through manual installation or using the provided Docker image to increase data privacy and accessible computational resources.

## 5 Discussion

cOmicsArt is a GUI designed to leverage interactive visualization during exploratory analysis while ensuring a seamless transition to further analysis by guaranteeing GUI-independent reproducibility. While cOmicsArt shares some functionalities with existing web-based GUIs, it has – to the best of our knowledge – its unique position in the landscape of tools: recognizing the collaborative nature of research in the life sciences, cOmicsArt prioritizes simplicity in initial phases of exploration and intentionally avoids overwhelming users with excessive options at the interface itself, yet still allowing for customization through code modification. The HTML report feature provides a clear overview of the analysis, providing a single point of reference for documentation and discussions.

While the modular backend allows for easy additions to the analysis suite, a key limitation is that updates to the R code snippets provided to users require manual implementation. Any new analysis must be integrated into both the Shiny modular framework and the downloadable R scripts, as changes in cOmicArt’s analysis are not automatically reflected in the user-accessible code. Furthermore, we plan to expand the outreach efforts for cOmicsArt to students and the biomedical society, enhance integration with public databases and repositories, and include support for multi-omics data analysis. Our long-term goal is to empower a broader community to work efficiently and empower collective strengths in biomedical research.

## Acknowledgements

We sincerely thank all early users of cOmicsArt for their invaluable feedback and patience while working with an app during its development. We also thank our students for participating so eagerly during the introduction of cOmicsArt.

