## Supplementary for "cOmicsArt – a customizable Omics Analysis and reporting tool"

**Supplementary Figures**
**Figure S1** Screenshot of a cOmicsArt module.
**Figure S2** Required data input.

**Supplementary Tables**
**Table S1** Comparison of available bulk-omics analysis tools with a graphical user interface
**Table S2** All major R packages and respective references.

**Supplementary Note A**
Description of systematic literature research to identify comparable tools to cOmicsArt

**Supplementary Note B**
Comparison to Galaxy and BioExpress

**Supplementary Note C**
Utilizing cOmicsArt for Iterative Data Analysis: From Data Input to Comprehensive Insights and Reproducibility

**Supplementary Note D**
Moving from cOmicsArt to R: Customizing result visualisations and performing additional analyses

**Supplementary E**
HTML- report accompanying analysis of Supplementary Note C

**Supplementary F**
Zip-folder containing R code and data accompanying analysis of Supplementary Note D

**Supplementary References**

### Supplementary Figures

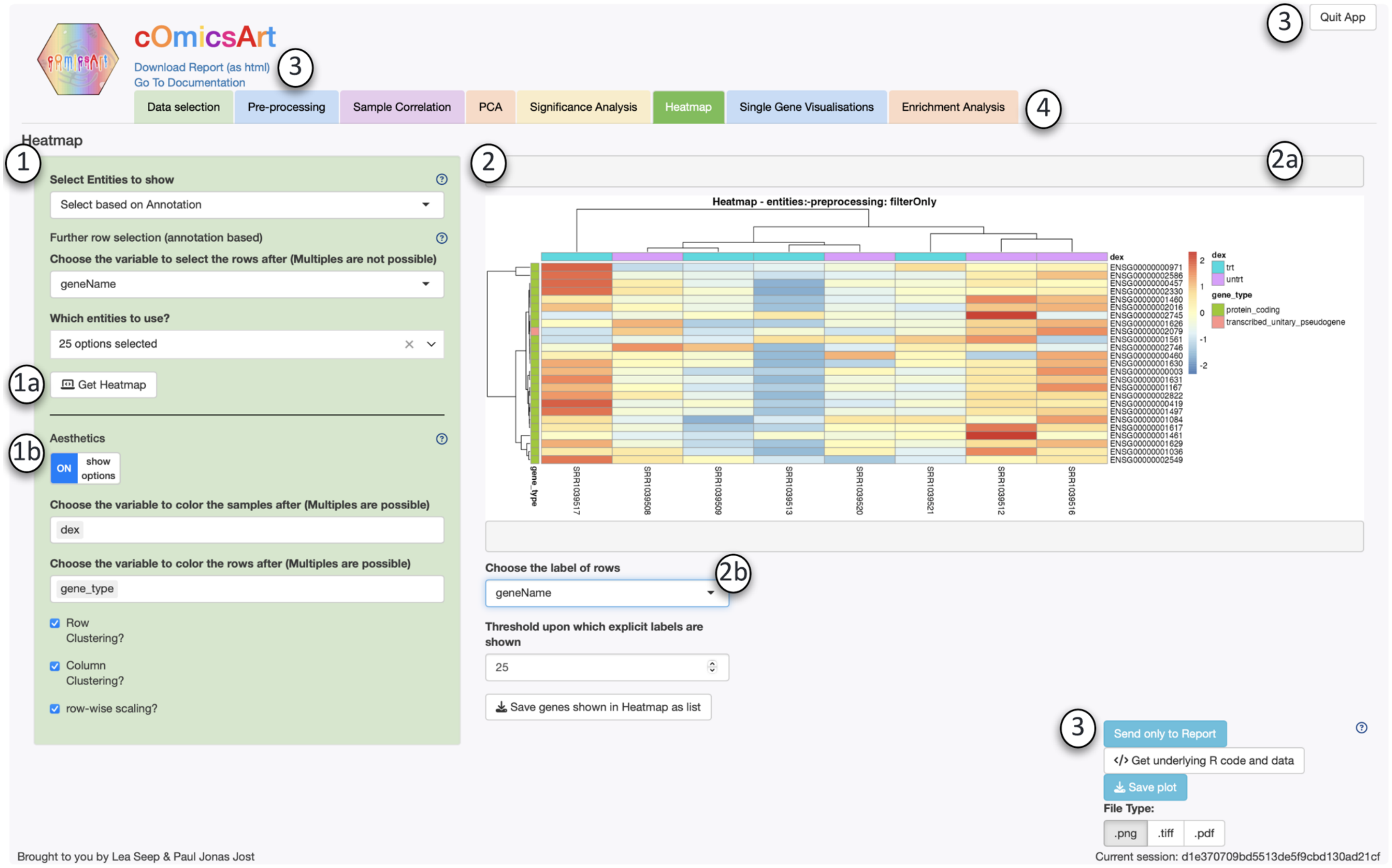


**Figure S1 Screenshot of a cOmicsArt module.** Every module is designed with the same design features to allow users to quickly use each panel once the design is understood. The recurrent structure includes the following: **1** The *Side panel* holds the parameters and options for the respective analysis. The sidebar is divided by a horizontal line into an upper and lower panel. **1a** The *upper panel* includes all options that require re-computation, triggered only when the 'Get ...' button is clicked. **1b** The *lower panel* includes options that immediately reflect changes in the displayed result. **2** The *Main panel* holds the analysis results (sometimes combining multiple mainpanel-tabs – not shown). **2a** The *notifications area* provides useful information or warnings to the user regarding the conducted analysis. **2b** The *display options & analysis-specific downloads* section offers direct customization options and additional analysis-result specific options. **3** The *Constant elements* are always visible to the user and include several download options, the link to the user documentation, and the quit button. The little blue question marks provide quick help for the respective element they are next to. **4** All accessible *tabs* shown. The user can switch between them by clicking onto them.

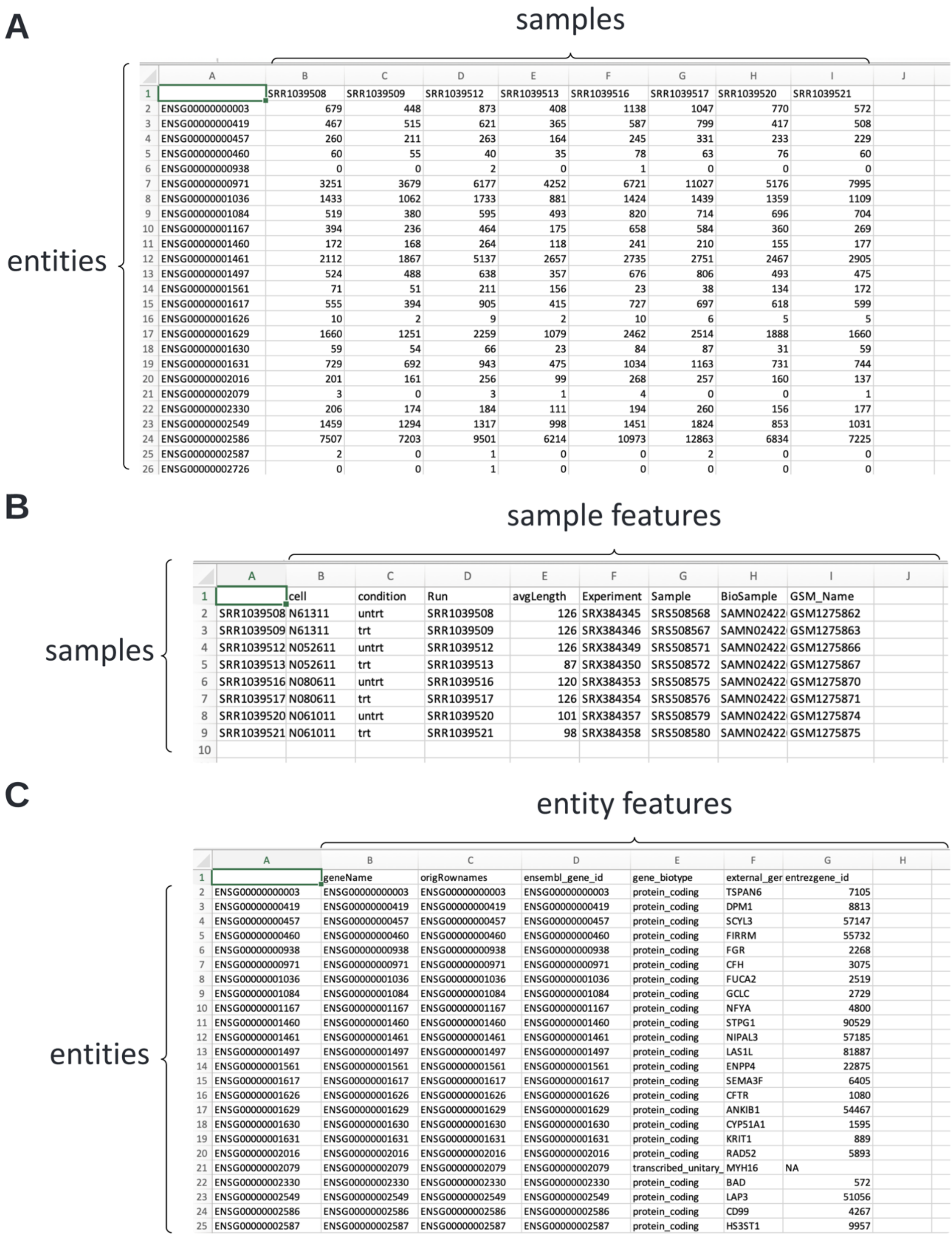


|  | A | B | C | D | E | F | G | H | I | J |
|---|---|---|---|---|---|---|---|---|---|---|
| 1 |  | SRR1039508 | SRR1039509 | SRR1039512 | SRR1039513 | SRR1039516 | SRR1039517 | SRR1039520 | SRR1039521 |  |
| 2 | ENSG00000000003 | 679 | 448 | 873 | 408 | 1138 | 1047 | 770 | 572 |  |
| 3 | ENSG00000000419 | 467 | 515 | 621 | 365 | 587 | 799 | 417 | 508 |  |
| 4 | ENSG00000000457 | 260 | 211 | 263 | 164 | 245 | 331 | 233 | 229 |  |
| 5 | ENSG00000000460 | 60 | 55 | 40 | 35 | 78 | 63 | 76 | 60 |  |
| 6 | ENSG00000000938 | 0 | 0 | 2 | 0 | 1 | 0 | 0 | 0 |  |
| 7 | ENSG00000000971 | 3251 | 3679 | 6177 | 4252 | 6721 | 11027 | 5176 | 7995 |  |
| 8 | ENSG00000001036 | 1433 | 1062 | 1733 | 881 | 1424 | 1439 | 1359 | 1109 |  |
| 9 | ENSG00000001084 | 519 | 380 | 595 | 493 | 820 | 714 | 696 | 704 |  |
| 10 | ENSG00000001167 | 394 | 236 | 464 | 175 | 658 | 584 | 360 | 269 |  |
| 11 | ENSG00000001460 | 172 | 168 | 264 | 118 | 241 | 210 | 155 | 177 |  |
| 12 | ENSG00000001461 | 2112 | 1867 | 5137 | 2657 | 2735 | 2751 | 2467 | 2905 |  |
| 13 | ENSG00000001497 | 524 | 488 | 638 | 357 | 676 | 806 | 493 | 475 |  |
| 14 | ENSG00000001561 | 71 | 51 | 211 | 156 | 23 | 38 | 134 | 172 |  |
| 15 | ENSG00000001617 | 555 | 394 | 905 | 415 | 727 | 697 | 618 | 599 |  |
| 16 | ENSG00000001626 | 10 | 2 | 9 | 2 | 10 | 6 | 5 | 5 |  |
| 17 | ENSG00000001629 | 1660 | 1251 | 2259 | 1079 | 2462 | 2514 | 1888 | 1660 |  |
| 18 | ENSG00000001630 | 59 | 54 | 66 | 23 | 84 | 87 | 31 | 59 |  |
| 19 | ENSG00000001631 | 729 | 692 | 943 | 475 | 1034 | 1163 | 731 | 744 |  |
| 20 | ENSG00000002016 | 201 | 161 | 256 | 99 | 268 | 257 | 160 | 137 |  |
| 21 | ENSG00000002079 | 3 | 0 | 3 | 1 | 4 | 0 | 0 | 1 |  |
| 22 | ENSG00000002330 | 206 | 174 | 184 | 111 | 194 | 260 | 156 | 177 |  |
| 23 | ENSG00000002549 | 1459 | 1294 | 1317 | 998 | 1451 | 1824 | 853 | 1031 |  |
| 24 | ENSG00000002586 | 7507 | 7203 | 9501 | 6214 | 10973 | 12863 | 6834 | 7225 |  |
| 25 | ENSG00000002587 | 2 | 0 | 1 | 0 | 0 | 2 | 0 | 0 |  |
| 26 | ENSG00000002726 | 0 | 0 | 1 | 0 | 0 | 0 | 0 | 0 |  |

|  | A | B | C | D | E | F | G | H | I | J |
|---|---|---|---|---|---|---|---|---|---|---|
| 1 |  | cell | condition | Run | avgLength | Experiment | Sample | BioSample | GSM_Name |  |
| 2 | SRR1039508 | N61311 | untrt | SRR1039508 | 126 | SRX384345 | SRS508568 | SAMN02422 | GSM1275862 |  |
| 3 | SRR1039509 | N61311 | trt | SRR1039509 | 126 | SRX384346 | SRS508567 | SAMN02422 | GSM1275863 |  |
| 4 | SRR1039512 | N052611 | untrt | SRR1039512 | 126 | SRX384349 | SRS508571 | SAMN02422 | GSM1275866 |  |
| 5 | SRR1039513 | N052611 | trt | SRR1039513 | 87 | SRX384350 | SRS508572 | SAMN02422 | GSM1275867 |  |
| 6 | SRR1039516 | N080611 | untrt | SRR1039516 | 120 | SRX384353 | SRS508575 | SAMN02422 | GSM1275870 |  |
| 7 | SRR1039517 | N080611 | trt | SRR1039517 | 126 | SRX384354 | SRS508576 | SAMN02422 | GSM1275871 |  |
| 8 | SRR1039520 | N061011 | untrt | SRR1039520 | 101 | SRX384357 | SRS508579 | SAMN02422 | GSM1275874 |  |
| 9 | SRR1039521 | N061011 | trt | SRR1039521 | 98 | SRX384358 | SRS508580 | SAMN02422 | GSM1275875 |  |
| 10 |  |  |  |  |  |  |  |  |  |  |

|  | A | B | C | D | E | F | G | H |
|---|---|---|---|---|---|---|---|---|
| 1 |  | geneName | origRownames | ensembl_gene_id | gene_biotype | external_ger | entrezgene_id |  |
| 2 | ENSG00000000003 | ENSG00000000003 | ENSG00000000003 | ENSG00000000003 | protein_coding | TSPAN6 | 7105 |  |
| 3 | ENSG00000000419 | ENSG00000000419 | ENSG00000000419 | ENSG00000000419 | protein_coding | DPM1 | 8813 |  |
| 4 | ENSG00000000457 | ENSG00000000457 | ENSG00000000457 | ENSG00000000457 | protein_coding | SCYL3 | 57147 |  |
| 5 | ENSG00000000460 | ENSG00000000460 | ENSG00000000460 | ENSG00000000460 | protein_coding | FIRRM | 55732 |  |
| 6 | ENSG00000000938 | ENSG00000000938 | ENSG00000000938 | ENSG00000000938 | protein_coding | FGR | 2268 |  |
| 7 | ENSG00000000971 | ENSG00000000971 | ENSG00000000971 | ENSG00000000971 | protein_coding | CFH | 3075 |  |
| 8 | ENSG00000001036 | ENSG00000001036 | ENSG00000001036 | ENSG00000001036 | protein_coding | FUCA2 | 2519 |  |
| 9 | ENSG00000001084 | ENSG00000001084 | ENSG00000001084 | ENSG00000001084 | protein_coding | GCLC | 2729 |  |
| 10 | ENSG00000001167 | ENSG00000001167 | ENSG00000001167 | ENSG00000001167 | protein_coding | NFYA | 4800 |  |
| 11 | ENSG00000001460 | ENSG00000001460 | ENSG00000001460 | ENSG00000001460 | protein_coding | STPG1 | 90529 |  |
| 12 | ENSG00000001461 | ENSG00000001461 | ENSG00000001461 | ENSG00000001461 | protein_coding | NIPAL3 | 57185 |  |
| 13 | ENSG00000001497 | ENSG00000001497 | ENSG00000001497 | ENSG00000001497 | protein_coding | LAS1L | 81887 |  |
| 14 | ENSG00000001561 | ENSG00000001561 | ENSG00000001561 | ENSG00000001561 | protein_coding | ENPP4 | 22875 |  |
| 15 | ENSG00000001617 | ENSG00000001617 | ENSG00000001617 | ENSG00000001617 | protein_coding | SEMA3F | 6405 |  |
| 16 | ENSG00000001626 | ENSG00000001626 | ENSG00000001626 | ENSG00000001626 | protein_coding | CFTR | 1080 |  |
| 17 | ENSG00000001629 | ENSG00000001629 | ENSG00000001629 | ENSG00000001629 | protein_coding | ANKIB1 | 54467 |  |
| 18 | ENSG00000001630 | ENSG00000001630 | ENSG00000001630 | ENSG00000001630 | protein_coding | CYP51A1 | 1595 |  |
| 19 | ENSG00000001631 | ENSG00000001631 | ENSG00000001631 | ENSG00000001631 | protein_coding | KRIT1 | 889 |  |
| 20 | ENSG00000002016 | ENSG00000002016 | ENSG00000002016 | ENSG00000002016 | protein_coding | RAD52 | 5893 |  |
| 21 | ENSG00000002079 | ENSG00000002079 | ENSG00000002079 | ENSG00000002079 | transcribed_unitary_ | MYH16 | NA |  |
| 22 | ENSG00000002330 | ENSG00000002330 | ENSG00000002330 | ENSG00000002330 | protein_coding | BAD | 572 |  |
| 23 | ENSG00000002549 | ENSG00000002549 | ENSG00000002549 | ENSG00000002549 | protein_coding | LAP3 | 51056 |  |
| 24 | ENSG00000002586 | ENSG00000002586 | ENSG00000002586 | ENSG00000002586 | protein_coding | CD99 | 4267 |  |
| 25 | ENSG00000002587 | ENSG00000002587 | ENSG00000002587 | ENSG00000002587 | protein_coding | HS3ST1 | 9957 |  |

**Figure S2 Required data input.** cOmicsArt expects three data matrices. **A** The data matrix contains the entities, such as genes, lipids, or metabolites, in its rows with respective measurements for each sample across the columns. **B** Sample annotation matrix has the same samples in its rows, with each column occupied by a feature describing each sample, such as sample condition. **C** Row annotation matrix has the same entities in its rows, with each column occupied by a feature further describing each entity, such as alternative IDs or gene biotypes. Note that even if there is no additional information for the annotation table, at least one column matrix must be provided. Note, that the sample table can be replaced by a Metadatasheet.

### Supplementary Tables

| Tool | Required data Input | Analysis features | Data Processing | Supported Omic types | active | Last maintained | Multi-omics | AI-extracted highlights |
|---|---|---|---|---|---|---|---|---|
| cOmicsArt | Measurement matrix, sample and row annotation or rds object, or metadatasheet | DE analysis, Volcano Plots, Correlation, Heatmap, clustering, Enrichment, PCA, set analysis | sample & entitiy filtering, scaling, transformation, normalisation, DESeq processing, batch correction | Transcriptomics Lipidomics Metabolomics | ✓ | July 2024 | ✗ | 1. Seamless Integration 2. Reproducibility 3. Interactive Visualizations |
| Argonaut[1] | One file containing measurement and additional sample/row annotation. | Outlier detection, Volcano Plots, Scatter Plots, Correlations, PCA, Go Enrichment | filtering,log2-transform, imputation | All if in data matrix format | ✓ | 2022 | ✗ | 1. Code-Free Platform 2. Real-Time Statistical Analysis 3. Interactive Data Visualization |
| asterics[2] | Measurement Matrix | Heatmap, PCA, Self Organizing Maps, MFA, DE analysis | sample & entitiy filtering, scaling, transformation, normailzation, EdgeR, batch correction | Not tailored to omic type | ✓ | July 2024 | ✓ | 1.Exploratory and Integration Analysis 2. Interactive Plots 3. Comprehensive Graphical Workflow |
| BioJuppies[3] | FastQ files or Count Table in tabular format, | PCA, Clustering, DE analysis, Enrichments | Alignment | Transcriptomics | ✓ | 2022 | ✗ | 1. Automated Jupyter Notebooks 2. Interactive Data Visualization 3. Persistent Cloud Storage |
| ExpressVis[4] | Measurment matrix, sample annotation | DE analysis, Volcano Plots, Heatmaps, Enichment Analysis | based on omics type | Transcriptomics, Proteomics, Microarray data | ✓ | 2022 | ✓ | |
| ggVolcanoR[5] | LFCs and p-values | Volcano plots, Correlation, Heatmaps, Upsets plots | ✗ | Transcriptomics Proteomics | ✓ | 2022 | ✓ | 1. Customizable Data Visualization 2. User-Friendly Interactive Interface 3. Optimised Publication-Quality Plots |
| ideal[6] | Measurement matrix, sample annotation and (optinal) row annotation, or rds object | DE analysis, Enrichment analysis, | sample exclusion, DESeq processing, heatmap, clustering | Transcriptomics | ✓ | May 2024 | ✗ | 1.Reproducible Analysis 2. Interactive Web Application 3.Publication-Ready Visualizations |
| iDEP[7] | Measurement Matrix, Sample annotation | Heatmap, PCA, DE analysis, Enrichment, Volcano Plots | vst, rlog, EdgeR | Transcriptomics | ✓ | June 2024 | ✗ | 1.Differential Expression Analysis 2.Pathway Analysis 3. Intuitive User Interface |
| IRIS-EDA[8] | Measurement matrix, sample matrix, (if scRNAseq row matrix with bp-length of gene) | Correlations Heatmaps, clustering, PCA, MDS, t-SNE, coexpression network analysis | regularized (based on library size) log, log2-transform, vst | Transcriptomics (bulk & single cell) | ✗ | March 2021 | ✗ | 1.User-Friendly Platform 2.Comprehensive Visualization Tools 3.FAIR Data Principles |

Continued on next page

| | | | | | | | | |
|---|---|---|---|---|---|---|---|---|
| MetaboAnalyst[9] | various, depending on planned analysis | Statistical analysis (including PCA, Correlation Analysis, Volcano Plots, DE analysis), Dose response, Biomarkers, Enrichment analysis, network analysis, Causal Analysis | scaling, transformation, normalisation, spectral processing | Metabolomics | ✓ | June 2024 | ✗ | 1. Automated Parameter Optimization<br>2. Functional Meta-Analysis<br>3. Streamlined Data Analysis |
| MicroScope[10] | Measurement matrix | DE analysis, Heatmap, PCA, GO enrichment, Network analysis | ✗<br>done, but no choices | Transcriptomics, ChIP-Seq | ✓ | 2017 | ✗ | 1. Interactive Heatmap Production<br>2. Integrated Analysis Tool<br>3. User-Friendly Interface |
| normSeq[11] | Measurement matrix, sample annotation | DE analysis, Heatmap, PCA | filtering, normalization, batch correction | Transcriptomics | ✓ | Oct 2023 | ✗ | 1. Data Normalization<br>2.Information Gain<br>3. User-Friendly Platform |
| OmicsPlayground[12] | Measurement matrix, sample and row annotation,<br>contrast tables | DE analysis, descriptive statistics, set analysis, correlation, clustering, heatmap, PCA, tSNE, GSEA, functional analysis, signature, biomarker analysis, single-cell profiling | filtering, log2 transformation, quanti. Normalisation, batch correction | Transcriptomics<br>Proteomics<br>(bulk & singel cell) | (✓)<br>requires subscription if one wants to use online | July 2024 | ✗ | 1. User-Friendly Platform<br>2. In-Depth Analysis and Visualization<br>3. Single-Cell Data Analysis |
| O-Miner [13,14] | CEL files<br>IDAT files<br>GEO series number | DE analysis, Cytoband<br>conserved TFBS, Survival plot<br>Correlation, Copy neutral LOH<br>CN from genome seuqencing, CpG island level, Enrichments, Venn diagram | normalization, filtering, phenotype suggestion | Transcriptomic<br>Genomics<br>Methylomic | ✗ | n.a. (no source code deposit) | (✓)<br>muliple projects from TCGA | 1. High-throughput profiling<br>2. Bioinformatics analysis and annotation<br>3. Integration of heterogeneous data |
| PiMP [15] | MzXML file | Peak annotation, PCA,<br>statistical analysis, Volcano plots, total Ion chromatograms,<br>single entitie chromatograms, Mapping to pathways, Network analysis | alignment, batch correction, identification | Metabolomic | (✗)<br>no registration possible | June 2017 | ✗ | 1. Automated analysis<br>2. Biological interpretation<br>3. Evidence cards |
| POMAShiny [16] | Measurement matrix, sample annotation | Volcano plot, density plot , heatmap, PCA , cluster analysis,<br>advanced statistics (Univariate, multivariate, correlations, ML, GLM, permutations tests and more) | sample selection, imputation, normalisation, outlier detection | Metabolomics<br>Proteomics | ✗<br>(website) | May 2023 (last commit) | ✗ | 1. Biomarker discovery<br>2. Statistical analysis<br>3. User-friendly workflow |
| Shiny-Seq [17] | Measurement matrix, sample & row annotation | DE-analysis, Coexpression network analysis , GSEA,<br>TF binding site ORA | normalisation, batch correction | Transcriptomics | (✓)<br>paid registrati/ docker image broken | April 2019 | ✗ | 1. Comprehensive RNA-Seq data analysis<br>2. Batch effect estimation and removal<br>3. Enrichment analysis |
| ShinyOmics[18] | none<br>(data and results placed at the backend) | Plotting of results (e.g. DE analysis, change in fitness of all genes vs select gene)<br>comparison of results | ✗ | Tn-Seq,<br>Transcriptomics<br>Proteomics | ✓ * | December 2020 | (✓)<br>comparison of results possible | 1. Rapid collaborative exploration<br>2. Visualization and comparison options<br>3. Data management and online sharing |

Continued on next page

| | | | | | | | | |
|---|---|---|---|---|---|---|---|---|
| STAGEs [19] | RNAseq count matrix, Log2-normalised Data Ratios and p-values | volcano plots, DE-analysis, enrichment analysis (Gene Set and Pathway), cluster analysis, correlation | ✗ | Transcriptomics | ✓ ** | June 2024 | ✗ | 1. Interactive visualization<br>2. Pathway enrichment analysis<br>3. User-friendly customization |
| TCC-GUI [20] | tag count matrix or option to simulate data within gui | DE-analysis, PCA, Volcano Plots | robust normalizazion (based on TCC-method[21]) | Transcriptomics ChIP-Seq (tag count data) | ✓ | March 2024 | ✗ | 1. Robust normalization<br>2. Graphical user interface<br>3. Visualization tools |
| Visual Omics [22] | Measurement matrix, “input group file”, FASTA (for protein domain prediction), other tab separated analysis results (e.g. DESeq output), ggplot2 object | DE-analysis, enrichment analysis protein domain prediction, protein-protein interaction, PCA/PcoA, boxplots, heatmaps, set interaction, bubble charts, Volcano plots | scaling normalisation log10-transform | Transcriptomics Proteomics | ✓ | June 2023 (no github) | ✗ | 1. Scientific chart editing<br>2. Multiple omics analyses<br>3. One-stop data analysis |
| WilsON[23] | CLARION file (data type agnostic) | Boxplots, bar plots, line plots, violin plots, heatmaps, scatter plots, PCA, correlation | ✗ | all types of omics (open file format) | ✓ | June 2022 | ✗ | 1. Dynamic visualization<br>2. Interactive workbench<br>3. Multi-omics data analysis |

not under original website location, but: https://shinyappstore.com/a/shinyOmics

*Website not the one mentioned in the publication but on github (https://kohcl17-stages-mirror-stages-zvg7qu.streamlit.app/ )

**Table S1 Comparison of available bulk-omics analysis tools with a graphical user interface (set of characteristics 1/2)**. ✓ and ✗ indicated whether a feature is present/absent or true/false. (✓) hints to restrictions specified in the respective cell. ( ✗ ) means not given / unable to test (e.g. paid registration would have been needed). To obtain the AI-extracted highlights we used ChatGPT4o with the following prompt with the respective abstract: given the following abstract of a paper, give three buzzwords of the paper in the context of benefits in the Omics Analysis. Abbreviations: ORA = over-representation-analysis, DE = Differential expression, GSEA = gene set enrichment analysis, sc = single cell

| Tool | interaction with repositorys | Export options of results | Reproducibility within GUI | Reproducibility independent of GUI | Customisation options |
|---|---|---|---|---|---|
| cOmicsArt | ✗ | Plots (png, tiff, pdf), tabular results (csv, xlsx), Report (html), R Code and Data (zip) | ✓ | ✓ | ✓ |
| Argonaut[1] | ✗ | Plots (svg), Data within plots in tabular format (txt) | ✗ | ✗ | ✗ |
| asterics[2] | ✗ | Plots (png), Report (html, not reproducible) | ✗ | ✗ | ✓ |
| BioJuppies[3] | ✓ | Downloadable Jupyter Notebook which also contains figures. Allows separate download of figures in png Format | ✓<br>Jupyter Notebooks are created, which also serve as a report | ✓<br>Jupyter Notebooks are independent | ✓<br>multiple selection options within the app. Additionally Jupyter Notebooks can be adjusted |
| ExpressVis[4] | ✗ | Plots (png), tabular results (txt) | ✗ | ✗ | ✓ |
| ggVolcanoR[5] | ✗ | pdf, tabular results | ✗ | ✗ | ✓<br>adjustment of figure export witdth, report adjustmens |
| ideal[6] | ✗ | Png, pdf, jpeg, tabular results , html or md report, rds objects | ✗ | ✓ | ✓ |
| iDEP[7] | (✓ )<br>option for download Public Data | Plots (pdf, png, svg), tabular results (csv), Report (html), Partial R Code (R) | (✓ )<br>Report for each section available but not working | (✓ )<br>R Code for DE Analysis | ✓<br>multiple algorithms for Enrichment and wide variety of sets |
| IRIS-EDA[8] | (✓ )<br>Provides support for submission to GEO | Pdf, png, tabular results | ✗ | ✗ | ✗ |
| MetaboAnalyst[9] | ✓ | Png, pdf, tiff, sag, PostScript | ✗ | ✓ | ✓ |
| MicroScope[10] | ✗ | Plots (html), tabular results (csv) | ✗ | ✗ | ✓ |
| normSeq[11] | ✓ | Plots(png), tabular results (txt) | ✓<br>Parameters in json file | ✗ | ✗ |
| OmicsPlayground[12] | (✓ )<br>Count data from GEO | tabular results, png, pdf | ✗ | ✗ | ✓<br>e.g. how many labeled genes to display |
| O-Miner [13,14] | ✓ | Results downloadbale as text, graphic output and excel (stated not tested) | ( ✗ ) | ( ✗ ) | ( ✗ ) |
| PiMP [15] | ✗ | Pimp-specific xml files | ( ✗ ) | ( ✗ ) | |
| POMAShiny [16] | ✗ | tabular results, pdf report of Explorative data analysis, plots as png | ✗ | ✗ | ✗ |

Continued on next page

| | | | | | |
|---|---|---|---|---|---|
| Shiny-Seq [17] | ✗ | outputs to a powerPoint, tabular results | ✗ | ✗ | |
| ShinyOmics[18] | ✗ | Plots (png, svg, pdf), tabular results | ✗ | ✗ | ✓<br>brush - select genes which information is then displayed within the table |
| STAGEs [19] | ✗ | DEG and pathway analysis (xlsx), calculated charts collectively in one pdf-report | ✓<br>Report incorporates chosen parameters | ✗ | ✓<br>Set the colour bar range<br>adjust bar plot height |
| TCC-GUI [20] | ✗ | tabular results, report with option to select results to include / exclude | ✓<br>Report incorporates chosen parameters | ✓<br>DE-analysis script can be copied, partial code for visualisation given within report | ✓<br>Axis Label assignment + Title,<br>Hihglight color |
| Visual Omics [22] | ✗ | Tabular results (.csv), plots as png, ggplot objects as RData | ✗ | ✗ | ✓<br>proivdes 262 tuneable paramteres from ggplot2 series (a graph library for R), |
| WilsON[23] | ✗ | Plots (png, pdf) | ✓<br>log files keep record of all analysis steps (but are not descriptive in terms of set/ chosen parameters => this more hidden in Rdata object) or in separate json file | ✓<br>Rdata objects for manual reproduction and manipulation of the visualizations , also serve to encapsulate input and plot functions for long term storage | ✓<br>Change color scheme, type of plots can be chosen (line, boxplot etc.), scale of y axis can be chosen |

**Table S1 Comparison of available bulk-omics analysis tools with a graphical user interface (set of characteristics 2/2)**. ✓ and ✗ indicated whether feature is present/absent or true/false. (✓) hints to restrictions specified in the respective cell. ( ✗ ) means not given / unable to test (e.g. paid registration would have been needed). Reproducibility within GUI was counted as present if all user-choices were easily obtainable by the user for e.g. describing what exactly has been done. Abbreviations: ORA = over-representation-analysis, DE = Differential expression, GSEA = gene set enrichment analysis, sc = single cell

| Package | Ref. | Package | Ref. | Package | Ref |
|---|---|---|---|---|---|
| BiocManager | 24 | ggplot2 | 25 | stringr | 26 |
| clusterProfiler | 27 | SummarizedExperiment | 28 | tidyr | 29 |
| DESeq2 | 30 | readxl | 31 | viridis | 32 |
| Dplyr | 33 | AnnotationDbi | 34 | pcaPP | 35 |
| DT | 36 | biomaRt | 37 | RColorBrewer | 38 |
| grid | 39 | org.Hs.eg.db | 40 | ComplexUpset | 41 |
| pathview | 42 | org.Mm.eg.db | 43 | ggvenn | 44 |
| pheatmap | 45 | vsn | 46 | kableExtra | 47 |
| plotly | 48 | renv | 49 | gridExtra | 39 |
| shiny | 50 | devtools | 51 | UpSetR | 52 |
| shinyhelper | 53 | rsconnect | 54 | reshape2 | 55 |
| shinyjs | 56 | ReactomePA | 57 | svglite | 58 |
| shinymanager | 59 | knitr | 60 | purrr | 61 |
| shinyWidgets | 62 | zip | 63 | sva | 64 |
| ggplot2 | 25 | rmarkdown | 65 | testthat | 66 |

**Table S2 All major R packages and respective references.**

### Supplementary Note A – Description of systematic literature research to identify comparable tools to cOmicsArt

A systematic literature search was conducted with PubMed to identify relevant studies on graphical user interfaces (GUIs) and web-based tools for omics analysis. The initial prompt included terms like Graphical User Interface OR web-based OR platform and bulk-omics. A refined search was conducted incorporating user-friendly terms to identify relevant interfaces with a comparable target user as aimed for with cOmicsArt. Then a word count was performed on all abstracts, and ChatGPT4o was used to identify and extract all items associated with specific diseases or organisms, such as C. elegans. This process allowed us to systematically exclude studies related to particular diseases or organisms. The final PubMed prompt was:

```
(
"Graphical User Interface"[Title/Abstract] OR
"web tool"[Title/Abstract] OR
"web-based"[Title/Abstract] OR
"webpage"[Title/Abstract] OR
"website"[Title/Abstract] OR
"web application"[Title/Abstract] OR
"web applications"[Title/Abstract] OR
("platform"[Title/Abstract] AND
       ("web"[Title/Abstract] OR
       "interactive"[Title/Abstract]))
) AND (
"bulk"[Title/Abstract] OR
"bulk-omics"[Title/Abstract] OR
"omics"[Title/Abstract] OR
"RNA-Seq"[Title/Abstract] OR
"Transcriptomics"[Title/Abstract] OR
"Metabolomics"[Title/Abstract] OR
"Lipidomics"[Title/Abstract]
) AND (
"guided"[Title/Abstract] OR
"guide"[Title/Abstract] OR
"user"[Title/Abstract] OR
"users"[Title/Abstract] OR
"user-friendly"[Title/Abstract] OR
"easy to use"[Title/Abstract]
) NOT (
"cancer"[Title/Abstract] OR
"disease"[Title/Abstract] OR
"diabetes"[Title/Abstract] OR
"virus"[Title/Abstract] OR
"strongyloides"[Title/Abstract] OR
"yersinia"[Title/Abstract] OR
"influenza"[Title/Abstract] OR
"hypertension"[Title/Abstract] OR
"listeria"[Title/Abstract] OR
"huntingtons"[Title/Abstract] OR
"adenocarcinoma"[Title/Abstract] OR
"pathogen"[Title/Abstract] OR
"malaria"[Title/Abstract] OR
"protozoan"[Title/Abstract]
) AND (
2014:2024[pdat]
)
```

Titles and PubMed's highlighted matched terms were screened for relevance, excluding not-yet filtered species-specific studies, resource-only studies, distinct analysis implementations, and dataset/analysis results published via a GUI. The following abstract screening excluded further non-fitting studies, such as single-cell tools, peak identification tools, GUI-building tools, and integration-only tools. Full-text assessments removed non-web app tools, multi-omics required tools, and tools specific to certain algorithms. During testing, tools with websites that were no longer accessible were excluded. Note, that we did not contact the corresponding authors for support.

The final set included the broad-range platforms Galaxy and BioExpress which are separately compared in Supplementary Note B.

**Overview literature search** (conducted 19.08.2024):

Records identified through initial pubmed searching (n = 1204)
↓
Records screened focusing on user-friendliness (n = 591)
↓
Records that allow broad usage (n = 410)
↓
Abstract screening for eligibility (n = 77)
↓
Full Text screening for eligibility and hence included (n = 21)

### Supplementary Note B

### Comparison to Galaxy and BioExpress

Besides narrow purpose graphical user interfaces, there also exist general purpose platforms – two of those, Galaxy [67,68] and BioExpress [69] were also found in our systematic literature search.
Galaxy is an open-source platform that enables non-specialist researchers to perform complex bioinformatics analyses and create custom workflows through a web interface. It supports a wide range of analyses and offers extensive documentation to facilitate reproducibility. Galaxy is accessible on public servers, which makes it a popular choice for collaborative projects across the world. BioExpress is a cloud-based bioinformatics platform optimized for handling and analyzing large-scale genomic data. BioExpress focuses on efficient data management and high-throughput processing capabilities. Both platforms aim to simplify the bioinformatics workflow for researchers. However, Galaxy emphasizes ease of use and accessibility for researchers without computational backgrounds, while BioExpress focuses on scalable cloud-based solutions and the ability to handle large volumes of data efficiently.
cOmicsArt is a standalone option, which aims to provide a gentler learning curve for new users or those with limited time to familiarize themselves with general purpose platforms. It uniquely facilitates the transition to highly specialized analyses that are not readily achievable with Galaxy or BioExpress, thanks to its ability to export R code.
We believe, that these features make cOmicsArt a valuable tool next to general purpose platforms.

### Supplementary Note C

**Utilizing cOmicsArt for Iterative Data Analysis: From Data Input to Comprehensive Insights and Reproducibility**

This showcase demonstrates how to use cOmicsArt to analyze and visualize data based on a specific dataset.

As background information, the dataset used in this showcase comes from a study that investigates how a high-salt diet (HSD) impacts immune defense against kidney infections, utilizing transcriptomics data to explore these effects. The study finds that an HSD worsens pyelonephritis in mice by impairing neutrophil antibacterial function. This is due to urea accumulation and disrupted circadian glucocorticoid rhythms, which impair neutrophil function. For further information and additional data, please see the original publication by Jobin et al.[70].

The showcase is divided into sections: 'The Data – Retrieval and preparation, Upload and Pre-processing', 'Investigating global patterns - Sample Correlation and PCA', 'Statistical Analysis', 'Intermediate Summary', 'Final Documentation' and 'Final Summary'. Each section provides insights from the analysis itself as well as how to obtain them using cOmicsArt.

<u>The Data - Retrieval and Preparation, Upload and Pre-processing</u>

The raw count data and sample information were retrieved from the European Nucleotide Archive while the row annotation was created to fulfill cOmicsArt's requirements. After uploading the data to the cOmicsArt web application, we requested additional gene annotations and performed pre-processing using the DESeq option, successfully generating homogeneous distributions across samples, as indicated by diagnostic plots.

Retrieval and Preparation:

The raw data (fastq files) and the sample information can be retrieved from the European Nucleotide Archive under the project PRJEB28204 (https://www.ebi.ac.uk/ena/browser/view/PRJEB28204). The data was aligned to the mm10 reference genome with the STAR Aligner.

Having the data in a count matrix format, we adjusted the corresponding sample and created the row annotation tables. The sample annotation table contains information about the samples, whereby we kept the information on the treatment, organism, cell type, and individual sample ID. We ensured that all samples listed in the column names of the count matrix also appear in the row names of the sample annotations. The required row annotation table containing information about the genes was not given but created. Here, we initially only had the gene Ensembl IDs taken from the count matrix. We ensured that all entities listed in the rownames of the count matrix also appear in the row names of the row annotation. As a single column is minimally required for the row annotation, we added a column named *'Ensembl_ID'*, exactly coping the row names. Note, that there is extensive help with additional details for formatting data correctly within the documentation.

Within this showcase, we will come back to this step as we gain information throughout which we can add to our annotation table. All information provided in this step can be utilized during the analysis to filter and group the data or perform statistical tests on it.

Upload:

Upon creation of the sample and row annotation table, we can upload the data to the cOmicsArt web app. We select the omic type as *transcriptomics* and then provide the data matrix, the sample annotation, and finally the row annotation. A quick inspection of the data upon clicking the *'inspect data'* button shows that the data was prepped correctly and we can proceed with the analysis. To do so, we click on *'Upload new data'* and the data is loaded into the app.

As a first step, we add some more information to our entities. Specifically, we request cOmicsArt to add gene annotation on the basis of Mouse genes (GRCm39), which adds the gene names as well as the biotype of the genes to our entity annotation. Since we have named our initial column of the row

annotation '*Ensembl_ID'*, cOmicsArt can automatically detect the correct column to use as the basis to add the additional info within the provided table. The automatic detection needs to be confirmed by the user. Note, that you need to have either Ensembl IDs, Entrez IDs, or gene names for the addition of information to work. Upon clicking the *'Add Gene Annotation'* button, the data is added, before returning to the Data Upload tab to proceed.
For now, no selection of the data is done, such that all data is considered during the analysis, making us ready to *'Start the Journey'* by clicking on the respective button.

Pre-processing:
For pre-processing, we select the *DESeq* option. Upon selection, we specify *'Treatment'* as the main factor for the model. For more details about the DESeq2 pipeline used in the background, check out the DESeq2 vignette. The pre-processing involves filtering lowly expressed genes before applying the DESeq2 pipeline.

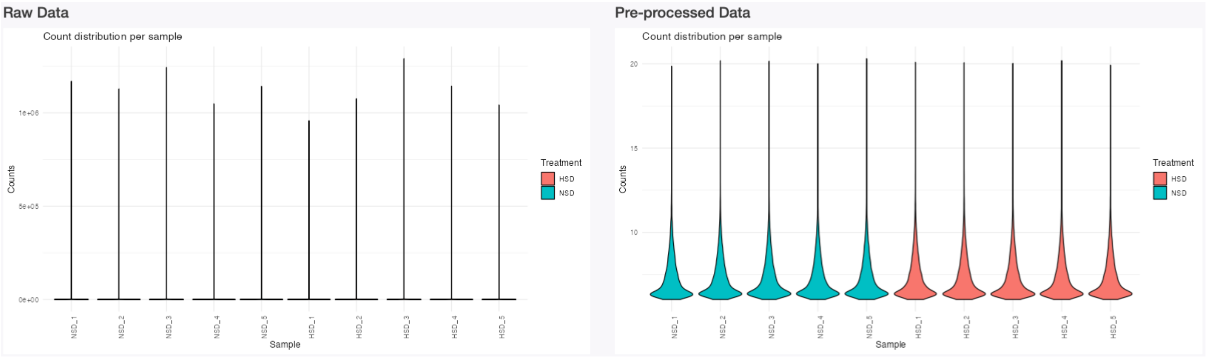

**Figure D1 Diagnostic plots**. The diagnostic plot displays each sample's gene count distribution colored after treatment.

After clicking *'Get Pre-Processing'*, the data is processed and we can assess the diagnostic plots of the sample distribution (Fig. D1). We color the sample distribution diagnostic plots with respect to their treatment (side panel below the horizontal line, hence only performing re-coloring but no re-pre-processing).
After preprocessing, we can see that the distribution across samples is quite similar and does not depend on the underlying treatment. We can conclude that the chosen pre-processing was successful in generating homogeneous distributions across samples.

Investigating global patterns - Sample Correlation and PCA
This section explores the correlation pattern among samples to determine if samples with similar treatments are more alike, utilizing sample correlation and principle component analysis.

Sample Correlation:
We first investigate whether global patterns within the data fit our expectations: samples with similar treatment are more similar to each other. Therefore, we assess their correlation pattern using the Pearson correlation coefficient within the sample correlation tab. Upon clicking *'Get Sample Correlation'* we can observe that the correlation across all our samples is very high. Upon selecting '*treatment'* as the annotation variable we can observe rather minor differences between the treatment groups (Fig. D2). This could indicate that the treatment does not have a strong effect on the global gene expression patterns. The expected effect is potentially only visible in a subset of the samples. Also, sample correlation incorporating all data might not be the best way to assess the effect of the treatment on such high-dimensional data. Hence, we further assess the PCA, to identify which linear combinations of features (here genes) explain the most variance within the data.

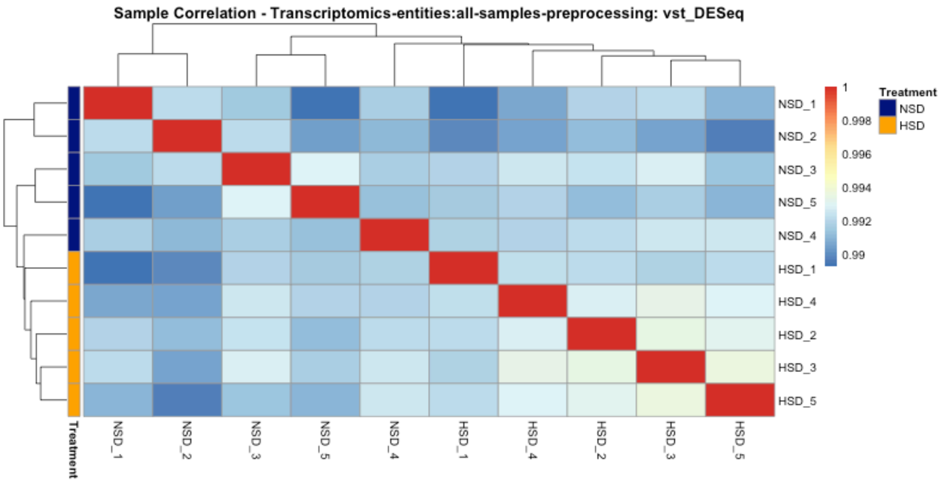


**Figure D2 Sample correlation plot.** The sample correlation matrix is displayed as a heatmap whereby hierarchical clustering is added and determines the ordering of rows/columns. Note, the small range of correlation values at the legend.

PCA:

Switching to the *PCA-panel* and performing a PCA colored after *Treatment*, we can observe that samples treated by HSD are less spread in the PC1 vs PC2 plot than the NSD samples (Fig. D3 A). Additionally, the samples have a separation tendency, although no clear separation is visible. We can also see that along PC1 (hence 'from left to right') two samples stand out on the left. Leveraging the tooltip (and changing *'Select the anno to be shown at the tooltip'* to *'individual_ID'*) we can identify the samples NSD_1 and NSD_2 as distinct from the other samples along PC1.

Notable is the rather low numbers of observed variance (indicated on the respective axis). A quick look at the scree plot tab confirms that the first three PCs explain roughly 18%, 15%, and 12% of the variance, respectively (Fig. D3 B). The following PCs all explain roughly the same amount of variance. This indicates that the data is rather complex and cannot be easily explained by a few linear combinations (PCA). Moroever, the NSD samples seem to be quite diverse which can be explained by the known functional diversity of neutrophil populations [71].

Switching to the *'PCA_Loadings'* tab allows us to gain insights into the features with the highest loadings for the principal component displayed on the x-axis. Here, we can choose underneath the plot *annotype shown on the y-axis*. This information is taken from our supplied row annotation, which was extended by our added gene annotation. When choosing the now available *gene_biotype*, we can see that among the highest loadings are genes of the type 'protein_coding'. Two non-protein coding genes are also among the highest loadings (TEC = to be experimentally confirmed and lncRNA, long-non-coding RNA) as well as three IDs (ENSMUSG00000095891, ENSMUSG00000075015, ENSMUSG00000075014) which do not have a biotype assigned. Within cOmicsArt such annotation NA's are replaced by the respective row ID. Switching the annotation to *external_gene_name*, we can see that Dusp1 and Ppbp are genes with the highest absolute loadings (Fig. D3 C). A quick literature search reveals that Dusp1 is a gene that has been identified to play a role in renal fibrosis, maintaining mitochondrial function and as a target of glucocorticoid-mediated signaling [72]. Ppbp (CXCL7) is a gene that has been identified as a chemoattractant for neutrophils[73].

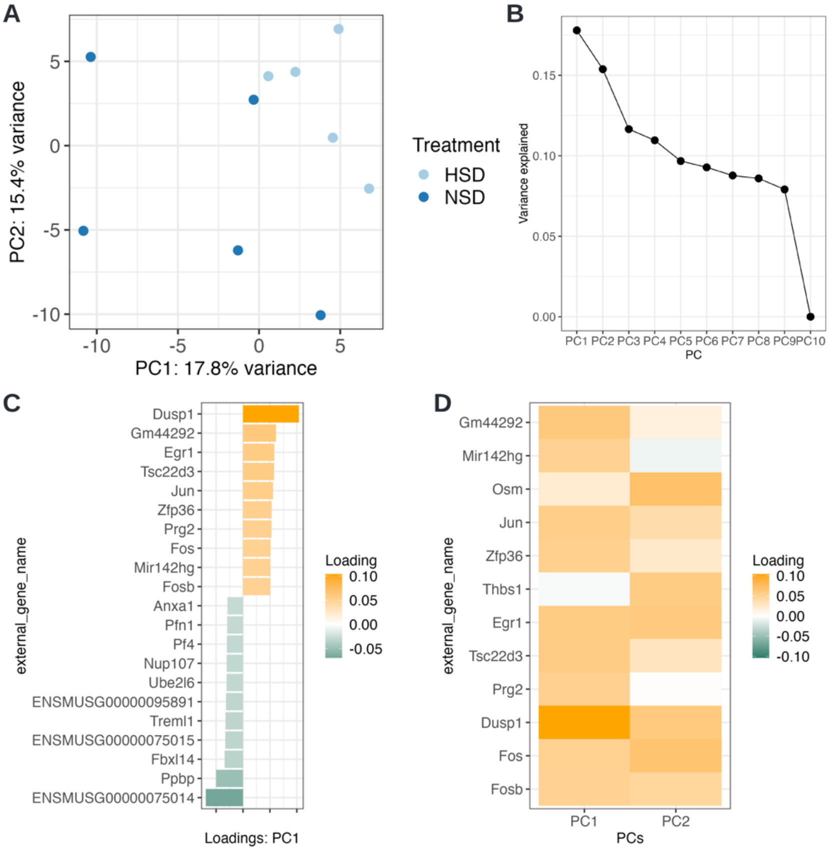


**Figure D3 PCA tab results.** A PCA plot showing 1st and 2nd principal components colored after treatment. B Scree-plot showing the variances explained per PC component. Within cOmicsArt one can hover over the points to retrieve the exact value. C The loadingsplot shows the top 20 absolute loadings and associated genes. Note that NA's in the intitial annotation are replaced by the respective row ID. D The loadings matrix plot shows the loadings greater than 0.05 for the selected PCs 1 and 2.

However, we can observe no separation of the groups along PC1 but rather by a combination of PC1 and PC2 (separation by a diagonal from middle top to bottom right) - hence high loadings on both principal components need to be assessed. For this, we switch to the *'PCA_Loadings_Matrix'* panel. To see something meaningful, we need to adjust the default parameters at the bottom. From the previous loadings plot, we can deduce that the general loading levels are rather small - therefore, we adjust the *'absolute loading threshold to filter entities with low impact'* to 0.05. As we are interested in PC1 and PC2, we adjust the number of PCs to include as well (Fig. D3 D). We can confirm by visual inspection that Dusp1 has the combined highest impact, followed by Fos and Osm. Fos in combination with Jun are components of the AP-1 transcription factor complex and have been reported to play a role in inflammaging[74]. Osm is a gene that has been known to have inflammatory and anti-inflammatory effects[75].

Statistical Analysis:

Following single interesting genes - Single Gene Visualisations:

To statistically test the two genes identified by their high loadings on PC1 and their difference in expression between the treatments, we can go to the *Single Gene Visualisations* tab. Here, single tests are performed and no multiple testing correction is done, allowing for a quick lookup of genes of interest. Checking Dusp1, Fos, Osm, and Ppbp, we obtain significant ($p<0.05$) results for all but Osm, with Dusp1 and Fos being upregulated in HSD compared to NSD, while Ppbp is downregulated (Fig. D4A).

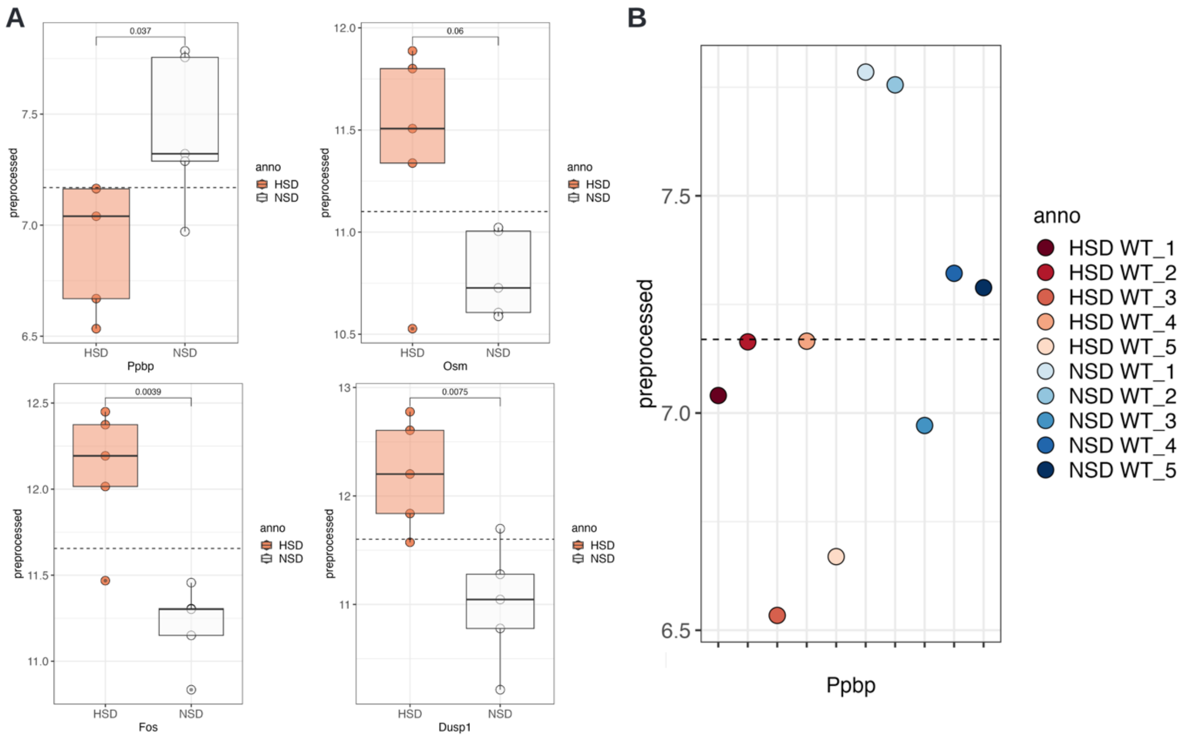


**Figure D4 Single gene visualisations.** A Boxplots with statistical testing for Ppbp, Osm, Fos, and Dusp1 expression grouped by treatment. B Dotplot of Ppbp expression shown per individual.

If we change the *'groups to show the data for'* to *'Simulation_Treatment'*, we can observe for Ppbp that NSD_1 and NSD_2 and HSD_3 and HSD_5 behave differently from the other members of the group (Fig. D4B). This aligns with the PCA results. Note, that we cannot 'test' for a difference as we have a single data point for each.

Assessing all genes – Significance analysis:
We switch to the *Significance analysis* tab to analyse all genes at once. We want to compare the treatment groups, specifically HSD vs. NSD, taking the latter as control. The order is important to interpret the direction of up- and down-regulation but does not change anything in terms of significance. As we have chosen the DESeq2 pipeline, cOmicsArt automatically selects the same test the pipeline. We obtain 47 genes (0.29% of the entire set) with significant changes between the conditions. The majority (31 genes) are significantly upregulated (16 downregulated) with a chosen significance level of 0.05 (after Benjamini-Hochberg multiple testing correction) (Table D1).
The most significant gene is ENSMUSG00000044786 (ZFP36). To get an overview of actual effect sizes (fold changes), we subselect within the shown table to show only the significant genes by clicking into the respective *padj* column in the table (where 'all' stands). Here we can adjust the sliding bar to select only genes with a padj value in the determined range. A quick check at the bottom of the table confirms we only selected the 49 entries. We then sort the log2Fold changes by clicking on the little grey up and down arrows. The Log2Fold change range goes from -0.34 to 1.23. Switching to the visual representation of the table, we go to the *tab Volcano*. Setting a Log FC threshold of 0.5, we can see that 10 genes remain as significant highlights (Fig. D5).

| Gene | baseMean | log2FoldChange | pvalue | padj |
|---|---|---|---|---|
| ENSMUSG00000024190 | 3425.50 | 1.23 | 2.69E-04 | 1.38E-02 |
| ENSMUSG00000038418 | 3400.16 | 1.06 | 1.21E-05 | 1.18E-03 |
| ENSMUSG00000021250 | 3324.20 | 0.95 | 3.25E-06 | 5.20E-04 |
| ENSMUSG00000052684 | 875.64 | 0.76 | 3.56E-05 | 2.34E-03 |
| ENSMUSG00000031431 | 1159.30 | 0.72 | 2.37E-04 | 1.32E-02 |
| ENSMUSG00000044786 | 8404.73 | 0.71 | 2.37E-19 | 2.65E-16 |
| ENSMUSG00000052837 | 8531.19 | 0.60 | 1.24E-07 | 2.48E-05 |
| ENSMUSG00000020423 | 1718.03 | 0.58 | 1.33E-07 | 2.48E-05 |
| ENSMUSG00000053560 | 1702.31 | 0.56 | 1.55E-08 | 5.78E-06 |
| ENSMUSG00000021123 | 655.70 | 0.52 | 8.26E-06 | 9.99E-04 |
| ENSMUSG00000042265 | 990.08 | 0.48 | 2.49E-05 | 1.99E-03 |
| ENSMUSG00000055148 | 1430.02 | 0.47 | 1.15E-07 | 2.48E-05 |
| ENSMUSG00000071076 | 1220.37 | 0.46 | 3.42E-10 | 1.92E-07 |
| ENSMUSG00000026034 | 1026.68 | 0.44 | 2.39E-05 | 1.99E-03 |
| ENSMUSG00000059657 | 599.05 | 0.43 | 6.41E-05 | 3.77E-03 |
| ENSMUSG00000021134 | 1140.64 | 0.41 | 2.96E-05 | 2.20E-03 |
| ENSMUSG00000030142 | 775.57 | 0.35 | 5.19E-04 | 2.15E-02 |
| ENSMUSG00000035569 | 2044.15 | 0.32 | 1.26E-05 | 1.18E-03 |
| ENSMUSG00000026987 | 2535.04 | 0.30 | 8.92E-06 | 9.99E-04 |
| ENSMUSG00000047888 | 1138.43 | 0.29 | 1.87E-03 | 4.56E-02 |
| ENSMUSG00000021025 | 5177.26 | 0.29 | 1.21E-03 | 3.56E-02 |
| ENSMUSG00000031229 | 1653.37 | 0.29 | 4.03E-06 | 5.63E-04 |
| ENSMUSG00000040054 | 749.37 | 0.29 | 5.97E-04 | 2.30E-02 |
| ENSMUSG00000041235 | 738.21 | 0.28 | 5.51E-04 | 2.20E-02 |
| ENSMUSG00000058318 | 823.56 | 0.26 | 1.71E-03 | 4.44E-02 |
| ENSMUSG00000022521 | 915.62 | 0.25 | 1.60E-03 | 4.26E-02 |
| ENSMUSG00000000078 | 4568.06 | 0.24 | 7.86E-04 | 2.66E-02 |
| ENSMUSG00000030083 | 1691.89 | 0.23 | 1.79E-03 | 4.44E-02 |
| ENSMUSG00000008348 | 8243.71 | 0.23 | 4.03E-04 | 1.81E-02 |
| ENSMUSG00000042406 | 2495.90 | 0.21 | 7.13E-04 | 2.57E-02 |
| ENSMUSG00000034994 | 2752.88 | 0.19 | 7.66E-04 | 2.66E-02 |
| ENSMUSG00000020846 | 4210.24 | -0.17 | 3.91E-04 | 1.81E-02 |
| ENSMUSG00000059182 | 2211.07 | -0.18 | 1.32E-03 | 3.68E-02 |
| ENSMUSG00000022584 | 1659.75 | -0.18 | 5.07E-04 | 2.15E-02 |
| ENSMUSG00000028249 | 2915.46 | -0.21 | 1.39E-03 | 3.79E-02 |
| ENSMUSG00000040659 | 1452.83 | -0.21 | 1.09E-03 | 3.29E-02 |
| ENSMUSG00000022372 | 1751.24 | -0.21 | 2.06E-03 | 4.89E-02 |
| ENSMUSG00000069516 | 41354.07 | -0.21 | 3.96E-04 | 1.81E-02 |
| ENSMUSG00000020849 | 1157.78 | -0.23 | 6.29E-04 | 2.35E-02 |
| ENSMUSG00000059108 | 2624.48 | -0.23 | 1.24E-03 | 3.57E-02 |
| ENSMUSG00000021537 | 773.58 | -0.24 | 8.65E-04 | 2.77E-02 |
| ENSMUSG00000024142 | 782.36 | -0.24 | 8.14E-04 | 2.68E-02 |
| ENSMUSG00000064246 | 585.36 | -0.26 | 9.44E-04 | 2.93E-02 |
| ENSMUSG00000033213 | 3082.27 | -0.26 | 4.47E-05 | 2.78E-03 |
| ENSMUSG00000029322 | 1406.33 | -0.27 | 2.72E-04 | 1.38E-02 |
| ENSMUSG00000019960 | 1965.95 | -0.30 | 1.77E-03 | 4.44E-02 |
| ENSMUSG00000102051 | 1772.11 | -0.34 | 3.15E-05 | 2.20E-03 |

**Table D1 Statistical analysis.** Overview showing all significant genes from the comparison HSD vs NSD (adjusted p value < 0.05 and sorted by log2FoldChange(LFC)). Omitting the columns 'lfcSE', baseMean and 'stat'. The double line separates up from downregulated genes.

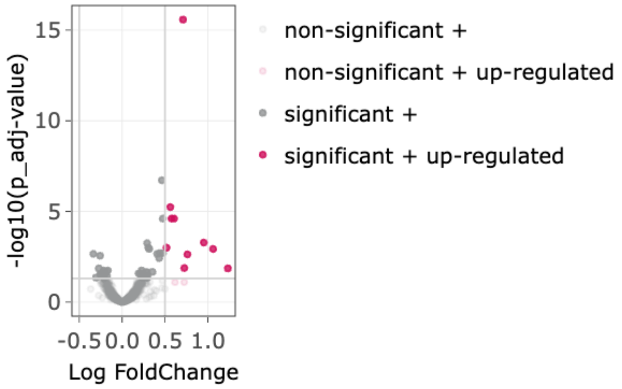


**Figure D5 Volcano plot.** Volcano plot of all tested genes, highlighting in red all signficant genes defined by adj. pvalue > 0.05 and a LFC > 0.5.

The set includes: ENSMUSG00000044786 (Zfp36), ENSMUSG00000052684 (Jun), ENSMUSG00000053560 (Ier2), ENSMUSG00000020423 (Btg2), ENSMUSG00000052837 (JunB), ENSMUSG00000021250 (Fos), ENSMUSG00000038418 (Egr1), ENSMUSG00000021123 (Rdh12), ENSMUSG00000031431 (TSC22D3), ENSMUSG00000024190 (Dusp1).

Half of these genes have been associated with major functions of neutrophils, such as Egr1 for microbial killing, chemoattractant priming, NETosis, Fos, Btg2, and Dusp2 for small pore migration, and JunB for NETosis71. Further, Jun, Ier2, JunB, Fos, and Egr1 belong to the early immediate response, while Zfp36, TSC22D3 and Dusp1 belong to the glucocorticoid targets.

We can conclude that besides no great global shifts, neutrophil core functions seem to be altered by High salt diet.

<u>Set analysis - Heatmap & Enrichment Analysis:</u>

To obtain a nice visual representation, we switch to the heatmap panel and select for the row-selection *Select based on Annotation* - which means that we can select data based on their row annotation hence for example precisely their ID. We select the 10 genes which we just identified to be signficiantly upregulated.

The resulting heatmap, after row-wise scaling, shows a distinct separation of the treatments, whereby the sample NSD_5 clusters closes to HSD_1, which itself clusters away from the other HSD samples (Fig. D6).

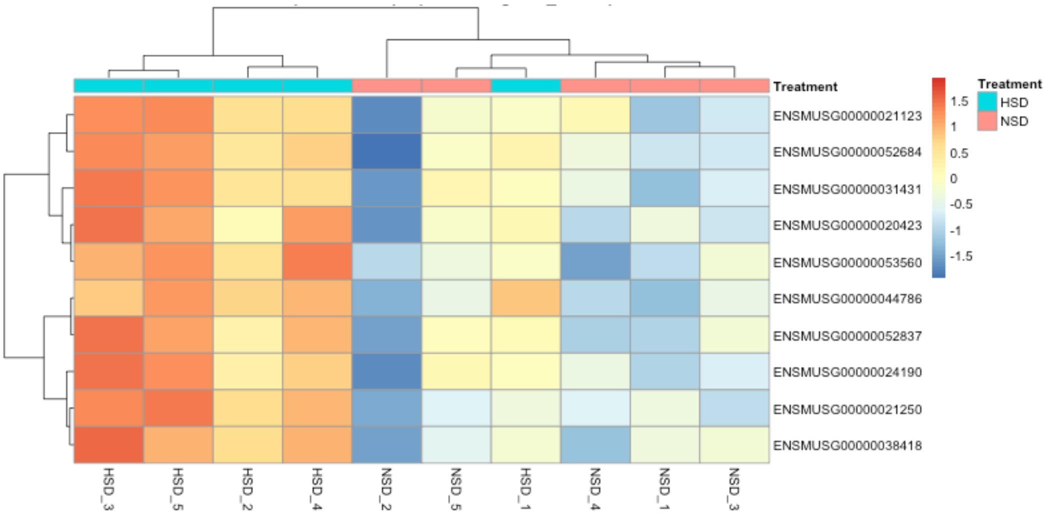


**Figure D6 Heatmap of significant upregulated genes.** The genes and the samples are ordered in correspondance to the applied hierarichal clustering.

We save the set of genes by clicking *Save genes shown in Heatmap for OA within Enrichment Analysis tab'* to further use them for overrepresantation analysis. To further characterise the set we can perform an enrichment analysis. We switch to the Enrichment Analysis tab and select the gene set we just identified. As we have a set of genes we first perform an Over-representation analysis uploading the identified set of genes. We choose the set to test as KEGG, HALLMARK and GO_BP (biological process). As universe, we used all genes present after pre-processing. We obtain enriched terms for Hallmark such as *TNFa signaling via NFkB*, *Hypoxia*, *UV response up,* and *P53 pathway* (Fig. D7).

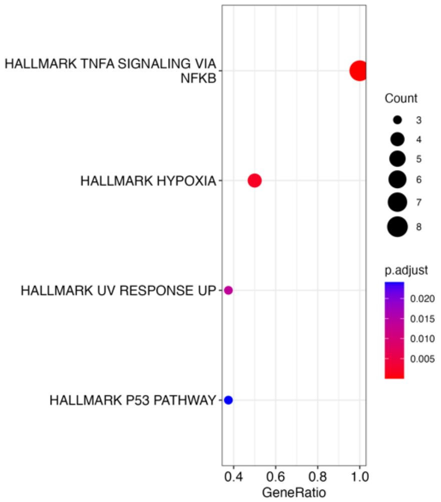


**Figure D7 Hallmark enrichment result.**

In addition to the visualisations we obtain the enrichment results as tables, telling us which query genes are within the respective (enriched) set. We can, for example, see, that 8 of our input genes belong to the term TNFa signalling via NFkB (Table D2).

| Description | GeneRatio | BgRatio | pvalue | p.adjust | geneID | Count |
|---|---|---|---|---|---|---|
| TNFA_SIGNALING_VIA_NFKB | 8/8 | 179/3573 | 3.41E-11 | 5.46E-10 | 12227/14281/19252/13653/ 22695/16476/16477/15936 | 8 |
| HYPOXIA | 4/8 | 163/3573 | 2.53E-04 | 2.03E-03 | 14281/19252/22695/16476 | 4 |
| UV_RESPONSE_UP | 3/8 | 135/3573 | 2.57E-03 | 1.37E-02 | 12227/14281/16477 | 3 |
| P53_PATHWAY | 3/8 | 182/3573 | 6.02E-03 | 2.41E-02 | 12227/14281/16476 | 3 |
| APOPTOSIS | 2/8 | 139/3573 | 3.61E-02 | 1.04E-01 | 12227/16476 | 2 |
| ESTROGEN_RESPONSE_LATE | 2/8 | 145/3573 | 3.90E-02 | 1.04E-01 | 14281/22695 | 2 |

**Table D2 Enriched Hallmark terms.** Only sets with at least two genes from the initial query set are shown.

Overall we observed rather subtle differences, for which it can be challenging to determine a set of interest that requires setting a threshold, e.g. LFC. To avoid setting such arbitrary threshold we performed a gene set enrichment analysis, which takes ranked entities into account, here chosen to be ranked by Log fold change (LFC). This approach is able to identify shifts of changes and is helpful if, for example, a single gene's change is not deemed significant or with a large effect size. The visualization of the enriched hallmark terms reveals the terms *MTORC1 signaling*, *Myc targets* and *Interferon alpha response*, as well as *Oxidative Phosphorylation*. The latter is also obtained for the KEGG set, next to *Alzheimer's*, *Huntington's*, and *Parkinson's disease*. GO also identified *Oxidative Phosphorylation* as an enriched term, along with *chromatin organisation*, *temperature homeostasis*, *positive regulation of acute inflammatory response*, and others (Fig. D8).
Considering the cell type (bone-marrow neutrophils), the studied cells seem to differ in their utilization of oxidative phosphorylation, which is a sign of differentiation for neutrophils[71]. Note that in the original publication[70], the ranking for GSEA was done based on t-test statistic values as well as a different DE-analysis was conducted as a basis; here it was done based on LFC values.

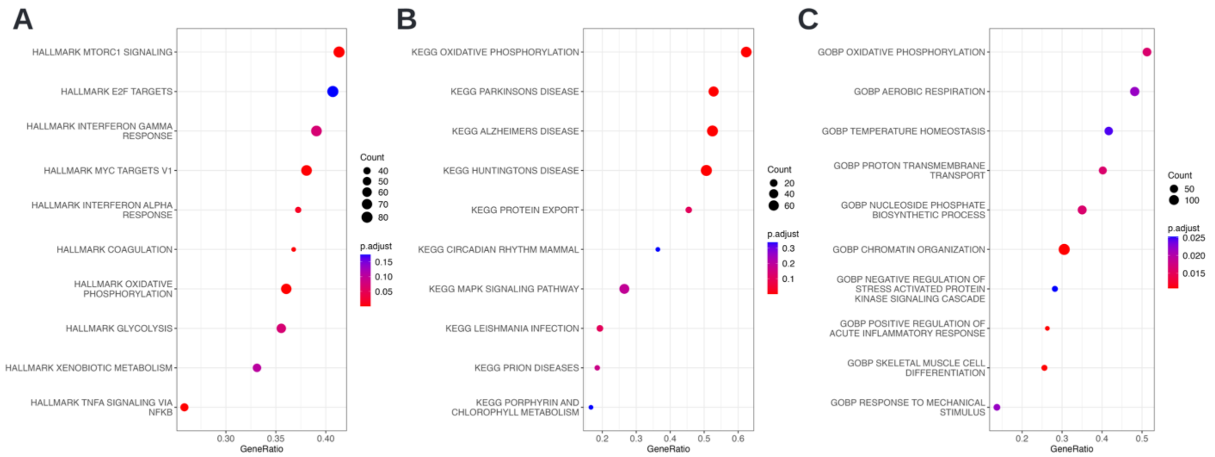


**Figure D8 Dotplots.** Enriched sets based on the Log Fold change between HSD and NSD. **A** Hallmark, **B** KEGG, **C** GO biological process.

<u>Intermediate summary</u>

We saw that on a global scale, we could not observe a clear pattern to distinguish between the treatments. This is, for example, globally visible when assessing sample correlation, as the correlation is overall at a high level. When looking at the PCA, we can see a rather high spread of samples belonging to NSD, whereas the HSD samples are less spread within the dimension reduction plot. Additionally, the statistical analysis returns only a small set of differentially expressed genes, indicating that the treatment effect affects a smaller portion of the entire data set. When performing an overrepresentation analysis of the DE-genes, we obtain a clear signal for TNFa signaling via NFkB. When performing gene set enrichment analysis on the LFC-ranked genes among the most enriched terms oxidative phosphorylation stands out. This together suggests that the effect of the HSD treatment alters the cellular metabolism in a directed fashion to an inflammatory state.

For further analysis, one might be interested in subselecting the data to focus on the potentially relevant aspects. For this, one can add the information from the statistical analysis to the gene annotation. This information is within the results table and can be obtained from the *significance analysis* tab. Moreover, one could add information to the entities indicating whether they are associated with the term *Oxidative phosphorylation* (using GO as a resource) to be able to visualize that specific subset within the *heatmap panel*. Additionally, while it may not be appropriate in this context, one could consider adding information to the samples, such as marking potential outliers and then redoing the analysis.

<u>Final Documentation</u>

All of the figures shown here were also sent to the report during the analysis of the dataset. The full HTML report, including the figures, can be found in Supplementary E. Some notes were added to showcase how the report can replace additional notes and be self-sufficient. If you explore the report, you can also find some publication-ready snippets to incorporate into your methods section.
Please note that while we've taken extensive measures to prevent data loss in the event of an unexpected crash of the web app, we cannot fully guarantee that it won't occur. To safeguard your work, we recommend periodically saving your report.
The analysis using cOmicsArt involved data retrieval, preparation, upload, and pre-processing, followed by a detailed investigation of sample correlation, PCA, and gene set enrichment to understand the subtle effects of HSD treatment in neutrophils.

### Supplementary Note D

#### Moving from cOmicsArt to R: Customizing result visualisations and performing additional analyses

This showcase demonstrates how to use cOmicsArt to customize visualizations to fit, e.g., a publication style. Additionally, the obtained analysis script is adjusted to highlight the option of customizing the analysis itself.
For context, the metabolomics dataset used in this showcase comes from a study investigating the effect of maternal obesity on the liver health state of the offspring[76]. The focus of the study is on Kupffer cells, liver-specific macrophages essential for liver development and functions. The findings reveal that maternal obesity disrupts Kupffer cell development in the fetus, leading to fatty liver disease and liver inflammation in the offspring in adulthood. Among other data, serum metabolomics was obtained under different conditions, precisely variations in the diet — high fat diet (HFD) or control diet (CD) — of the mother, of the foster mother, i.e. the lactating female, and of the offspring. Therefore a condition termed for example 'HFD CD CD' specifies that the mother recived HFD, while the lactating female, as well as the offspring CD. Serum metabolomics measurements were obtained from the offspring. When the mother received an HFD, this is referred to as maternal obese otherwise as maternal lean.
The showcase is divided into three sections: 'Data Upload and Pre-processing', 'PCA Analysis' and 'Additional Analysis'. The first section covers the transition from raw data to cOmicsArt and then to R, including basic customization. The second and third sections focus entirely on the analysis within R.

<u>The Data Upload and Pre-processing</u>

The data was received from Metabolon and prepared as detailed in the manuscript[76]. The obtained data sheets within an Excel workbook can be accessed from this repository (https://github.com/LeaSeep/MaternalObesity/tree/main/data – Metabolon.xlsx).
For uploading to cOmicsArt, the already pre-processed data matrix was transposed to fit the format accepted by cOmicsArt and then saved as a CSV file. Information about the metabolites, such as the pathways they belong to and additional identifiers, was placed in a separate row annotation table. Similarly, details about the samples were collected and organized into an independent sample annotation matrix for cOmicsArt. The original 'CHEM_IDs' were numerical, which could be mistaken with indexes when used as rownames. To resolve this, the prefix 'CHEM_ID' was added to all IDs within the data matrix and the row annotation table. Additionally, the hyphens ('-') within the sample names were replaced with underscores ('_'), as hyphens are not allowed as columnnames inside cOmicsArt.
The three data matrices were uploaded to cOmicsArt with no data selection performed. Also 'None' was chosen as pre-processing, due to the data pre-processing done outside the app. From the diagnostic plots colored after 'GROUP_NAME' we can observe that each distribution is centered around zero and a wide distribution of ranges between the samples. Due to the amount of samples, the plot is compressed. To obtain a decompressed plot one can receive the R code and data by pressing on the respective button. Upon executing the obtained script within RStudio, the violin plots are immediately obtained and shown in the plots viewer panel of RStudio. Within RStudio's export functionality, one can adjust the plotting area manually to enlarge the plot area and hence decompress the plot (Fig E1).

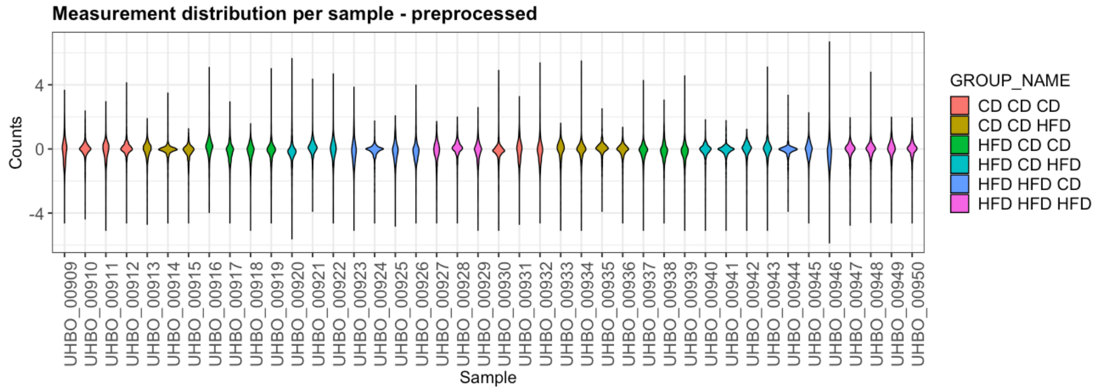


**Figure E1 Metabolomics measurement distribution per sample.** The plot is manually adjusted within Rstudio's plot panel to enlarge the plotting area and decompress the visualisation.

PCA Analysis

A PCA analysis is conducted to identify differences in the global metabolome between the maternal obese group and the maternal lean group. When the data is colored by 'GROUP_NAME', it becomes evident that the offspring's diet (hence the last diet out of the triplet) is the major contributing factor (Fig E2A). However, with the six groups displayed, distinguishing them is not straightforward. While one could define new groups, such as maternal lean v.s. maternal obese or offspring on CD v.s. HFD, to simplify the coloring, it is more useful to keep the groups separate and use similar colors to indicate similarities besides differing groups. This customization is not possible within the cOmicsArt user interface. However, users can obtain the R code and data to replicate the entire analysis, from data input to PCA visualization.

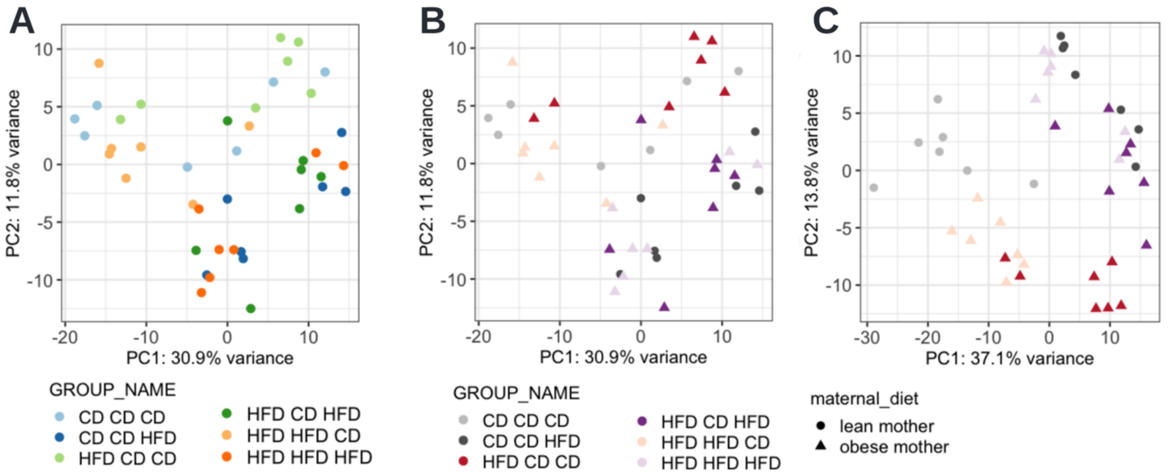


**Figure E2 PCA visualisation and analysis customisation. A** The original cOmicsArt generated PCA plot on batch-corrected data with default coloring. **B** Customised PCA-plot on the basis of obtained R script and data from cOmicsArt. The shape of the dots now resembles the maternal dietary status while the color theme was adjusted. **C** Customised batch correction, now supplying a model matrix, leads to a different appearing PCA result, leading still to the same conclusion.

The provided script allows for instant and self-sufficient reproducibility. By executing the script, the same PCA plot is obtained as a ggplot[25] object, which provides the basis for customisation. Here, an additional shape argument is given to indicate the mother's diet (CD vs HFD) and the color scheme was updated (Fig. E2B). Since ggplot is widely used, large language models like ChatGPT or GitHub Copilot can reliably assist in adjusting the code to meet specific needs.

One might note that we have not yet recreated the exact plot shown in the manuscript. The reason is that the two analyses differ in their batch correction procedure. When supplying a model matrix to the combat function, which biases the batch correction to retain as much variance as possible, which can be attributed to the variable of interest — in this case, 'GROUP_NAME', we could recreate the exact plot (Fig E2C) from the original publication. After these adjustments, the plot underscores that the most significant variance in the metabolomic data is due to the offspring's diet, rather than the mother's diet.

Additional analysis:

The straightforward approach to replicate cOmicsArt's results allow also for easy expansion of the analysis, through the reuse of code snippets or building upon supplied data in a convenient data object. We demonstrate how the PCA script can be reused for a different dataset. Then, the resulting PCA model is used to project left-out samples onto the constructed PCA space. Finally, we utilize the computed and projected principal components to quantify how predictive the offspring diet is within a machine-learning-based approach. Note that the complete code snippet – obtained from cOmicsArt and then expanded as described in the following – can be found in Supplementary.

The provided PCA code snippet can be re-used to compute and display a PCA with a different dataset. Here, we perform a PCA with a subset of mice, precisely those offspring that were born to mothers on CD, subsequently lactated by a foster mother on CD, and themselves received HFD or CD, respectively. The subsetted data object can either be used to overwrite the original data for direct reuse of the snippet or the code snippet needs to be adjusted to work with the new object.

The obtained PCA was then used to transform the left-out samples (i.e., those with differing maternal diets) to project them into the new PCA space. This approach allows us to visually assess how well the offspring's diet alone can separate samples with varying maternal diets. To avoid confusion, we added a 'p' to the group names of the projected samples. By adjusting the 'GROUP_NAME' column instead of adding a new column, we could simply re-use the previous plotting command. The resulting PCA (Fig. E3) shows that the offspring diet drives the separation, even the samples with differing maternal diets, indicating that the variance in the metabolome data is mainly attributed to the offspring's diet.

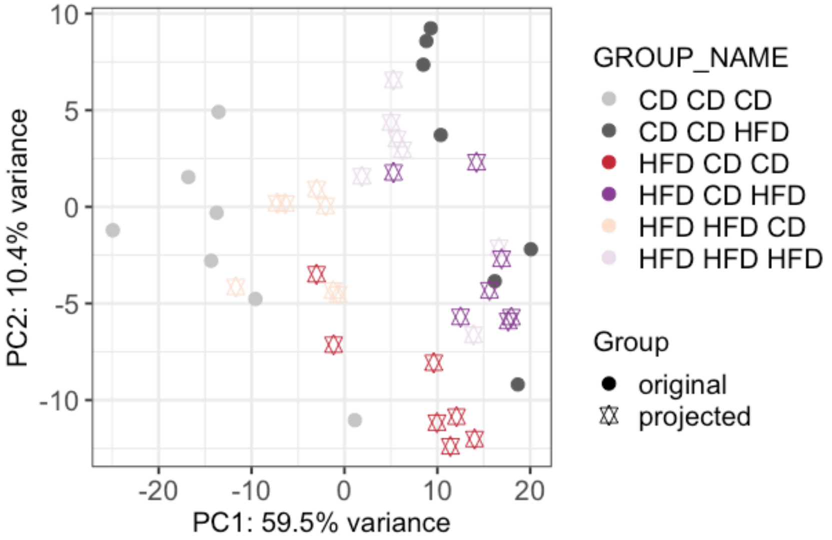


**Figure E3 Projection PCA.** Basis of PCA is built upon samples belonging to the group CD CD CD or CD CD HFD, while all other samples are projected onto that space, indicating how sufficient the offspring's diet is to separate the remaining samples according to their offspring diets regardless of their mother's diet.

While one can visually confirm that the offspring diet is the major source of variance within the metabolome data, we haven't quantified this. One possible quantification is by determining the predictive power of the offspring diet in predicting the offspring diet of the remaining samples with differing maternal diets. As descriptors for each sample, we use all the principal components from the above-described procedure (14 components). Due to the small sample size, we use k-nearest neighbors (k-NN) as the machine learning model, which does not require training but only the specification of the parameter k (the number of neighbors to consider), which is set to 3. Upon implementation, a prediction accuracy of 82% indicates a good prediction performance based solely on the offspring's diet. Assessing the true and false positive rates reveals a significant difference (0.6 vs. 0). The difference indicates that some samples with a true offspring CD are misclassified as having an offspring HFD (Table E1A). Further analysis of the respective maternal diets shows that the misclassified samples are from maternal obese, combined with an offspring diet of CD (Table E1B). Interestingly, this is not observed with an offspring diet of HFD. These findings highlight that while the offspring diet is the primary source of differences in the metabolome, the maternal diet, in combination with a CD offspring diet, induces detectable changes in the metabolome, making it more similar to that of an HFD offspring.

**A**

| Predicted \ Actual | CD | HFD |
|---|---|---|
| CD | 7 | 0 |
| HFD | 7 | 14 |

**B**

| Predicted \ Actual | HFD CD CD | HFD CD HFD | HFD HFD CD | HFD HFD HFD |
|---|---|---|---|---|
| CD | 2 | 0 | 5 | 0 |
| HFD | 5 | 7 | 2 | 7 |

**Table E1 Confusion matrices of kNN-based offspring-diet prediction. A** Confusion matrix indicating the correct classification of all HFD offspring samples, whereby seven additional actual CD samples have been wrongly predicted as HFD samples. **B** Confusion matrix showing the actual complete condition (of mothers and offspring) highlights that only samples with obese mothers on offspring kept on CD are misclassified as offspring on HFD.

The ability to access and customize code snippets directly from cOmicsArt significantly enhances the flexibility and depth of analysis for researchers. By providing the underlying R code, cOmicsArt not only allows for easy replication of visualizations and results but also allows users to extend and tailor their analyses to their needs and questions.
